# Identifying nonlinear Functional Connectivity with EEG/MEG using Nonlinear Time-Lagged Multidimensional Pattern Connectivity (nTL-MDPC)

**DOI:** 10.1101/2023.01.19.524690

**Authors:** Setareh Rahimi, Rebecca Jackson, Olaf Hauk

## Abstract

Investigating task- and stimulus-dependent connectivity is key to understanding how brain regions interact to perform complex cognitive processes. Most existing connectivity analysis methods reduce activity within brain regions to unidimensional measures, resulting in a loss of information. While recent studies have introduced new functional connectivity methods that exploit multidimensional information, i.e., pattern-to-pattern relationships across regions, they have so far mostly been applied to fMRI data and therefore lack temporal information. We recently developed Time-Lagged Multidimensional Pattern Connectivity for EEG/MEG data, which detects linear dependencies between patterns for pairs of brain regions and latencies in event-related experimental designs (Rahimi et al., 2022b). Due to the linearity of this method, it may miss important nonlinear relationships between activity patterns. Thus, we here introduce nonlinear Time-Lagged Multidimensional Pattern Connectivity (nTL-MDPC) as a novel bivariate functional connectivity metric for event-related EEG/MEG applications. nTL-MDPC describes how well patterns in ROI *X* at time point *t*_*x*_ can predict patterns of ROI *Y* at time point *t*_*y*_ using artificial neural networks (ANNs). We evaluated this method on simulated data as well as on an existing EEG/MEG dataset of semantic word processing, and compared it to its linear counterpart (TL-MDPC). We found that nTL-MDPC indeed detected nonlinear relationships more reliably than TL-MDPC in simulations with moderate to high numbers of trials. However, in real brain data the differences were subtle, with identification of some connections over greater time lags but no change in the connections identified. The simulations and EEG/MEG results demonstrate that differences between the two methods are not dramatic, i.e. the linear method can approximate linear and nonlinear dependencies well.

**Highlights:**

1. nTL-MDPC is a bivariate functional connectivity method for event-related EEG/MEG
2. nTL-MDPC detects linear and nonlinear connectivity at zero and non-zero lags
3. nTL-MDPC revealed connectivity between ATL hub and semantic control regions
4. Differences between linear and nonlinear TL-MDPC were small

## 1 Introduction

Cognitive processes originate from dynamic interactions among multiple brain regions (Bullmore and Sporns, 2009; Passingham et al., 2002). Connectivity methods are key to understand how brain regions interplay to generate these processes. The most common connectivity methods are unidimensional functional connectivity methods, which summarise information in a brain area by collapsing the pattern of activity across voxels or vertices. These methods may neglect valuable information in the brain response patterns (Anzellotti et al., 2017b, 2017a; Basti et al., 2019, 2018), as animal research has shown that inter-regional brain interactions are most likely multidimensional (DiCarlo et al., 2012). Thus, multidimensional connectivity methods have recently been proposed that make use of time courses for multiple voxels/vertices per region (Anzellotti and Coutanche, 2018; Basti et al., 2020; Rahimi et al., 2022b). Note that the terms “multivariate” and “multidimensional” on the one hand and “univariate” and “unidimensional” on the other have been used interchangeably in the field. As in our prior paper (Rahimi et al., 2022b) and similar to Basti et al. (2020), we here use “multidimensional” to refer to cases in which multiple time courses per brain region are explicitly considered. “Unidimensional” refers to scenarios in which a region’s time courses are summarised into one time course. Furthermore, in the computation of connectivity between two regions, we refer to “bivariate” and “multivariate” as cases where effects of two or multiple regions are being taken into account, respectively. These multidimensional connectivity methods are different from approaches that assess activity patterns per region but still estimate connectivity based on unidimensional time courses, e.g. using Representational Connectivity or Informational Connectivity Analysis (Karimi-Rouzbahani et al., 2022; Kriegeskorte et al., 2008; Laakso and Cottrell, 2000).

Recently, linear and nonlinear Multivariate Pattern Dependence (MVPD, NL-MVPD) (Anzellotti et al., 2017a, 2017b) were introduced to characterise statistical relationships between the multidimensional activity patterns of two ROIs in resting state functional magnetic resonance imaging (fMRI) data. In this approach, the dimensionalities of the data in each ROI are first reduced through principal component analysis (PCA). Then, the statistical relationship between the resulting factor loadings is tested either via linear regression or using a nonlinear artificial neural network (ANN). However, this dimensionality reduction prevents estimation of the original individual voxel-to-voxel relationships between patterns across regions. This issue has been addressed by Basti et al. (2019) for fMRI data, who used ridge regression to estimate the voxel-to-voxel transformations between pairs of ROIs and applied their method to event-related fMRI data. However, comparable methods for EEG/MEG data that can track multidimensional event-related connectivity over time are still emerging.

A few multidimensional connectivity approaches have been proposed specifically for electroencephalography and magnetoencephalography (EEG and MEG): the multivariate interaction measure (MIM, Ewald et al., 2012), multivariate lagged coherence (MVLagCoh, Pascual-Marqui, 2007), and multivariate phase-slope-index (MPSI, Basti et al., 2018). However, these are frequency-domain methods that require interpretation of effects in separate frequency bands. While some brain processes may indeed be reflected in specific frequency bands (Fries, 2015; Siegel et al., 2012), this does not necessarily hold for all brain processes, such as those reflected in early short-lived brain responses in event-related experimental paradigms. Also, the estimation of connectivity metrics in frequency bands results in a loss of temporal resolution, and does not allow the estimation of pattern relationships across different time lags. Thus, we require methods that estimate the time course of connectivity time sample-by-time sample.

Recently, Rahimi et al. (2022b) applied a similar approach to Basti et al. (2019) to EEG/MEG data in source space. Time-Lagged Multidimensional Pattern Connectivity (TL-MDPC) estimates the linear transformation between patterns for pairs of brain regions and pairs of time lags using cross-validated ridge regression. The explained variance (EV) of these linear transformations is used as a metric for connectivity strength. High EV suggests a strong relationship between pairs of ROIs across different time lags. EVs for different time lags for a given pair of ROIs can be presented in a time-time matrix, called a temporal transformation matrix (TTM) (a matrix plotting time in ROI X by time in ROI Y). The authors showed that TL-MDPC indeed captures multidimensional relationships between patterns that its unidimensional counterpart is insensitive to, in both simulated data and real EEG/MEG data contrasting two tasks with different semantic demands. TL-MDPC produced richer connectivity than unidimensional methods and distinguished between sub-networks reflecting meaningful divisions.

While the linear approach is computationally efficient and yields an easily interpretable transformation matrix, it may miss important nonlinear relationships between patterns. This is particularly important, as animal studies have shown that nonlinear relationships can be important (DiCarlo et al., 2012). Anzellotti et al. (2017b) showed that a nonlinear artificial neural network (ANN) identified more significant pattern dependencies between brain regions than a linear regression approach on resting-state fMRI data. Thus, in the present study we extended the linear approach of Rahimi et al. (2022) to create nonlinear Time-Lagged Multidimensional Pattern Connectivity (nTL-MDPC) by replacing ridge regression with an ANN as a method of finding the linear as well as nonlinear relationships between multidimensional patterns of activity in pairs of brain regions and across latencies.

Like TL-MDPC, nTL-MDPC is a bivariate undirected functional connectivity method suitable for event-related datasets. In contrast to similar methods that have previously been applied to fMRI data (as in Anzellotti et al., 2017a, 2017b; Basti et al., 2019), it characterises pattern connectivity over time, i.e. for different time lags, as well as over space. It does this by estimating how well patterns in ROI *X* at time point *t*_*x*_ can predict patterns in ROI *Y* at time point *t*_*y*_ through a nonlinear mapping. nTL-MDPC can estimate the full vertex-to-vertex transformations between ROIs. However, EEG/MEG source estimates have inherently limited spatial resolution which depends on source location, orientation, signal-to-noise ratio, etc. (Hauk et al., 2019; Molins et al., 2008; Samuelsson et al., 2021). Consequently, source signals in different voxels within an ROI carry redundant information (“leakage”). To address this issue, as in TL-MDPC, we employed a k-means clustering approach as a “feature selection” method to select the most informative vertices within the ROIs (Rahimi et al., 2022b). Unlike “feature extraction” methods such as PCA, here we do not transform features to create new ones, instead we simply select the most informative features (Khalid et al., 2014). We then estimate the nonlinear mappings between the sub-sampled patterns using a cross-validated ANN regression method, employing 10-fold cross validation and regularisation to avoid overfitting. As in TL-MDPC, the resulting cross-validated explained variance (EV) served as our connectivity metric.

As in our previous study, we assessed the performance of our model in both simulations and a real EEG/MEG dataset. In our simulations we tested to what degree nTL-MDPC is sensitive to 1) false positive errors for independent random patterns, 2) linear multidimensional patterns, and 3) nonlinear multidimensional patterns.. We evaluated the method for typical numbers of trials and vertices across a broad range of SNRs. In our real data analysis we contrasted dynamic semantic network connectivity between two word-based decision tasks with nTL-MDPC as we have previously done with coherence analysis (Rahimi et al., 2022a) and TL-MDPC (Rahimi et al., 2022b). Previously, we demonstrated limited connectivity between semantic ROIs when using spectral coherence as a unidimensional connectivity method (Rahimi et al., 2022a). TL-MDPC provided richer connectivity across a network of semantic representation and control regions. Here, we will use nTL-MPDC to test whether a nonlinear multidimensional connectivity method can corroborate and extend these findings.

## 2 Materials and Methods

We investigated whether the nonlinear nTL-MDPC method detects more connectivity than the linear TL-MDPC method in both simulated data and a real EEG/MEG dataset. We first introduce the rationale and procedures that are shared between both methods. In both cases, we first need to 1) prepare the patterns at each time in each region, 2) attempt to find a mapping/transformation between two patterns at two time points, 3) predict the target ROI’s output, 4) and finally measure the explained variance (EV) between the real and predicted output, as the connectivity metric.

Both methods have the same aim. Let us consider that ***X*** and ***Y*** are activity pattern matrices of ROI X and ROI Y at time point *t*_*x*_ and *t*_*y*_ of size *n*_*t*_ × *n*_*X*_ and *n*_*t*_ × *n*_*Y*_, respectively, where *n*_*t*_ is the number of trials, and *n*_*X*_ and *n*_*Y*_ are the number of vertices in the two regions. Figure 1a represents activity patterns in ROI X and ROI Y across time. We intend to find out whether there is an all-to-all mapping between the patterns of responses in the two ROIs at different latencies. Simply put, we are interested to see how well the patterns in ROI Y at time point *t*_*y*_ can be predicted from the patterns in ROI X at time point *t*_*x*_, and the other way around, through a transformation (in each direction).

**Figure 1.**
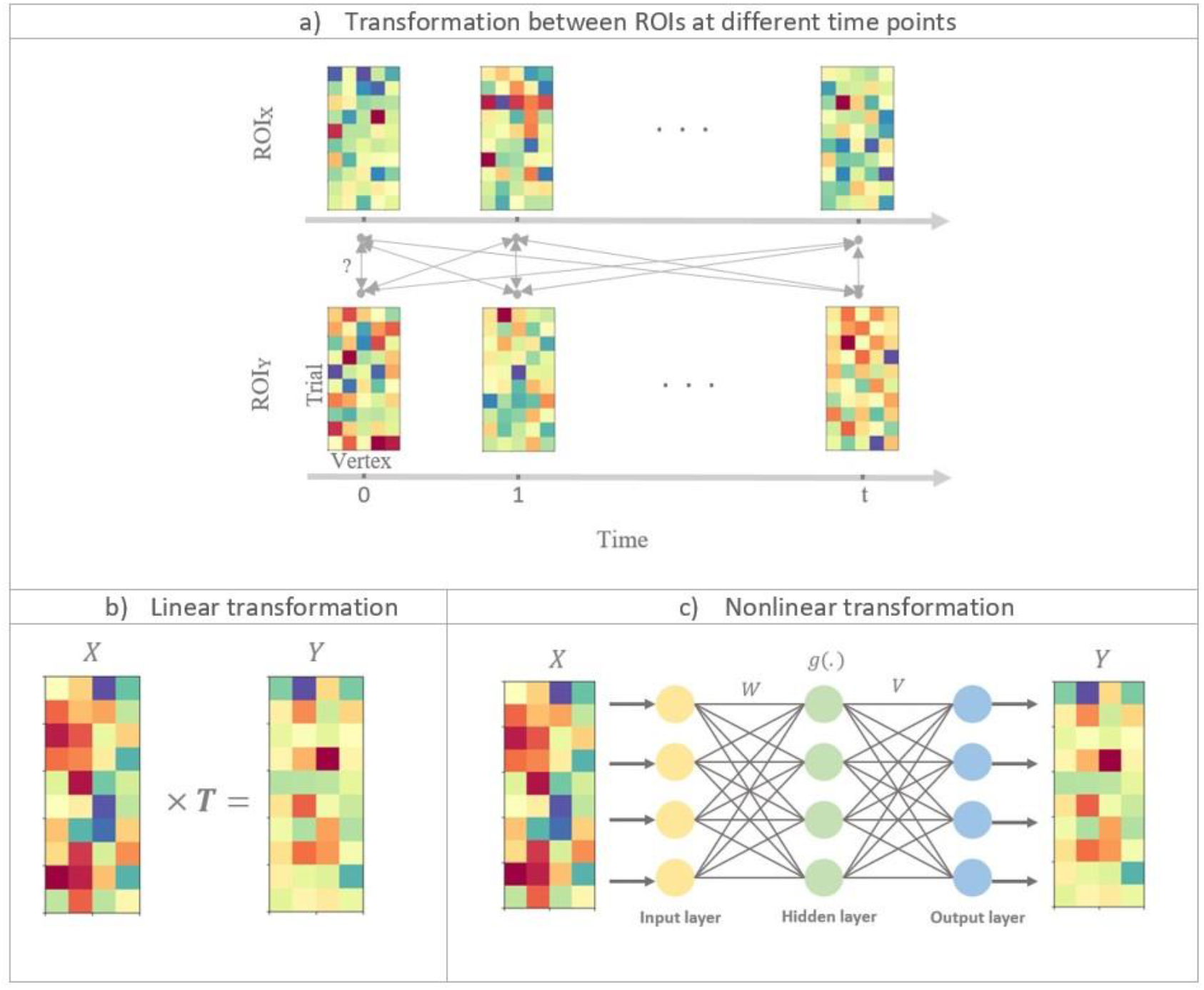
Illustration of linear and nonlinear time-lagged multidimensional pattern connectivity approaches. a) The principle of TL-MDPC is displayed. We assess the relationship between activity patterns in ROI X and ROI Y at different time lags. Each matrix indicates activity patterns in one ROI at one time point, with rows in each matrix indicating activation across different trials, and columns representing activation over different vertices in the ROI. Bidirectional arrows represent possible transformations and dependencies between patterns. b) Illustration of using TL-MDPC to detect the linear transformations ***T*** between patterns using ridge regression (as in Rahimi et al., 2022b). c) Illustration of the novel nTL-MDPC method to detect nonlinear (and linear) transformations between patterns using an artificial neural network.

To do so, we first need to address the spatial resolution of EEG/MEG. These signals are inherently smooth and have limited spatial resolution (Hauk et al., 2019; Palva et al., 2018). As a result, not all vertices are independent. To deal with this issue, unsupervised k-means clustering is employed as a “feature selection” approach, to sub-sample the most informative vertices within each ROI (Rahimi et al., 2022b). Using this approach, vertices serve as samples/observations and trials serve as features. Thus, we group all vertices into k clusters, so that all vertices with similar activation profiles across trials are within one cluster. To find out the optimum number of clusters, the elbow method (Ng, 2012) was used. There are several ways one could pick a representative vertex within each cluster, including computing the mean of all vertices, the centroid of the cluster, or the vertex with the highest variance. We pick the vertex with the highest variance as this allows us to keep the patterns in the genuine pattern space while estimating the transformations between regions. As a result of clustering, the patterns ***X*** and ***Y*** at time points *t*_*x*_ and *t*_*y*_ are now of size *n*_*t*_ × *n*_*x*_ and *n*_*t*_ × *n*_*y*_, where *n*_*x*_ and *n*_*y*_ are the number of clusters in ROI X and Y.

We are then able to estimate the transformations between regions. Using the dimensionality-reduced patterns in ROI X and Y at two specific time points (***X*** and ***Y*)** we can extract the transformation matrix ***T***, allowing us to predict the patterns in ROI Y 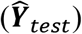 through *T* and test subset ***X***_*test*_. We can then measure the explained variance (EV) between the real output, ***Y***_*test*_ and the predicted output, and use this as a measure of connectivity. The highest score would be 1, suggesting a very strong relationship (either linear or nonlinear) between patterns, while close to zero values reflect very weak (or no) connectivity. In the following, we replace negative values of EV by zero since they indicate that the data could not fit by the model at all, and a variation in negative EVs is not meaningful. Additionally, due to the fact that our measurements are not directional, but a reflection of statistical dependencies between each pair, we predicted ***X*** from ***Y*** and ***Y*** from ***X***, and reported the average of resulting EVs.

For ROI X and Y, we repeat the whole procedure above for all possible pairs of time points, and the resulting EVs constitute the Temporal Transformation Matrix (TTM) for ROI pair X and Y. Thus, different areas in this matrix reflect connectivity at different time lags; the diagonal entries of TTMs show simultaneous connectivity between ROI X and Y patterns, the upper diagonal represents connectivity where *Y* is ahead, and lower diagonal shows connectivity where *X* is ahead.

### 2.1 The nTL-MDPC method

Although these methods share many steps, they differ in one critical aspect, the estimation of the transformations. While the linear TL-MDPC uses cross-validated regularised ridge regression (Hoerl and Kennard, 1970) (Figure 1b), the nonlinear method employs ANNs to extract the mapping between ***X*** and ***Y*** (Figure 1c). Artificial neural networks (ANNs) can estimate the relationship between linear and nonlinear multidimensional time courses (Singh et al., 2003). ANNs learn to map from a given input to a target output, in this case from the multidimensional activity patterns in one ROI at one time point to those in another ROI at one time point. As shown in Figure 1c, the activation in each unit in the ANN (other than input units) is generated through the weighted sum of the activity of connected units in the prior layer. In the hidden layer (i.e., the units which do not directly receive input or give output), unit activity is further determined with an additional step whereby every node passes the weighted sum of the inputs to that unit through a nonlinear function. In cases where the unknown coefficients outnumber the observation samples (an ill-posed or underdetermined problem), a regularisation procedure is needed. For this purpose, we used a cross-validated regularised ANN to avoid overfitting. As in the linear method, the explained variance of the transformations will serve as the connectivity metric.

#### 2.1.1 Organisation of the ANN

Figure 1c illustrates the general framework of a feedforward ANN (Venkatesan and Anitha, 2006) with one hidden layer. We used a nonlinear regression implemented in Python^1^ using ANN. We utilised a tangent hyperbolic function as the activation function as this is widely used (Sharma et al., 2017). The number of units in the hidden layer was set to be the average of the number of nodes in input and output.

The network is trained using a standard back-propagation learning algorithm (Rojas, 1996). The solver for weight optimisation was set to “LBFGS” (Limited-memory Broyden–Fletcher–Goldfarb–Shanno), an unconstrained nonlinear optimisation algorithm in the family of quasi-Newton methods. In general, LBFGS can have better performance than other available algorithms for small datasets^1^. Each pattern was standardised before being input into the ANN by subtracting the mean and dividing by the standard deviation of the pattern.

#### 2.1.2 Modelling statistical dependence using ANN

A general mapping between ***X*** and ***Y*** can be defined as follows:

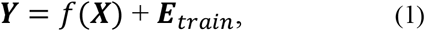

where *f*(·) could be a linear or nonlinear function, and ***E*** is a zero-mean Gaussian matrix. For nTL-MDPC, we use an ANN to have a more general estimation of *f*(·) than a linear regression allows. Using the train subset, for any ROIs X and Y, and a set of time points *t*_*x*_ and *t*_*y*_, we have:

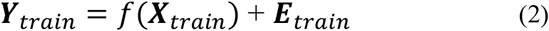

where ***X*** is the input of the ANN which is used to predict ***Y*** during training. Below, is a step-by-step description of how ***X*** and ***Y*** are related.

The input of the *h*th node in the hidden layer, *z*_*h*_, is obtained as follows:

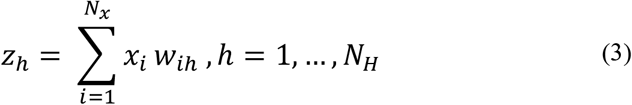

where *x*_*i*_ is the *i*th input, *w*_*ih*_ is the weight between the *i*th node in the input and the *h*th node in the hidden layer, *N*_*x*_ is the number of nodes in the input layer, and *N*_*H*_ is the number of nodes in the hidden layer.

The output of the *h*th node in the hidden layer, is as follows:

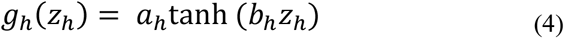

where *a*_*h*_ and *b*_*h*_ are scaling parameters of the tangent hyperbolic function. Finally, the output of the *j*th node in the output layer is obtained as:

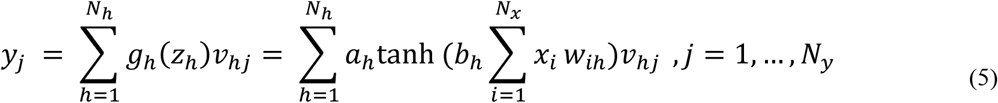

where *v*_*hj*_ is the weight between *h*th node in the hidden and *j*th node in the output layer, and *N*_*y*_ is the number of nodes in the output layer.

After estimating the above parameters and function *f*(·), the predicted patterns in ROI *Y* can be obtained using the test subset as follows:

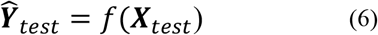

where 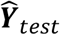 is the predicted pattern in ROI Y at time point *t*_*y*_. Explained variance (EV) is then computed for each vertex k=*1*,…, *n*_*y*_:

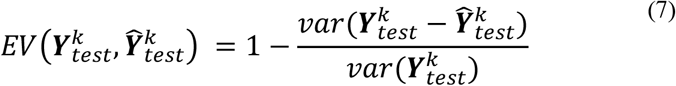

Finally, to quantify the multidimensional connectivity metric for each pair of ROIs at each pair of time points, we summarise the above EVs by averaging across all of the vertices in ROI Y:

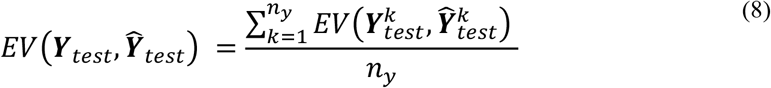

### 2.2 Simulations: Performance of linear and nonlinear TL-MDPC for different relationships between patterns

In this section, we compare the TL-MDPC and nTL-MDPC on simulated data for reasonable and practical choices of number of trials, vertices, and signal-to-noise ratios (SNR), to find out whether the nonlinear model can capture more connectivity than the linear one. TL-MDPC should be optimal when patterns are only linearly connected, i.e., when no nonlinearities are present. The nonlinear method should still be able to capture linear connectivity, but it is not clear whether this will be at the same level as the linear method and whether it can capture additional nonlinearities with the available training data. To this end, we created three different scenarios: 1) no relationship between the activity patterns, 2) linear multidimensional relationships and 3) nonlinear multidimensional relationships. As the estimation of transformations can be done time sample-by-time sample and does not rely on the precise time points of ***X*** and ***Y***, we here assess the performance of our method without simulating time courses. Thus, we use the shorter terms MDPC and nMDPC to refer to the two methods in this section. Figure 2a shows the first scenario, representing two independent patterns with no connectivity, so that there is no f(.). e.g., ***Y*** ≠ *f*(***X***).

**Figure 2.**
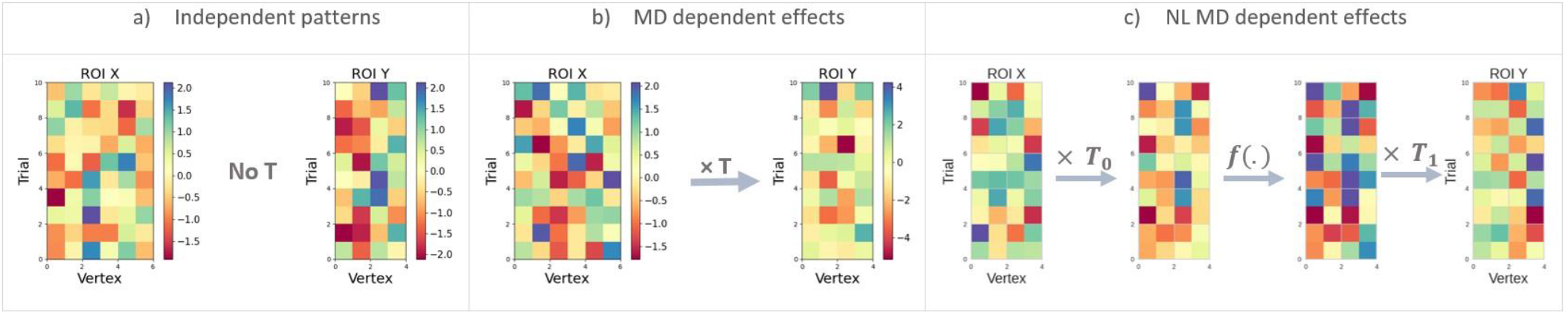
Representation of our simulation scenarios with different types of connectivity. The matrices (***X*** and ***Y***) show patterns of responses in ROI X and ROI Y, respectively, with rows indicating different trials and columns indicating vertices. a) ***X*** and ***Y*** are independent patterns with no reliable transformation between the regions and as a result no connectivity. b) There is a linear multidimensional relationship between ***X*** and ***Y*** through matrix ***T***. c) Nonlinear multidimensional relationships between ***X*** and ***Y*** are simulated through a neural network transformation. Patterns in ROI X are first being transformed through ***T***_0_ as a linear mapping, then the resulting output passed through a nonlinear (sigmoid or tangent hyperbolic) function *f*(·), and the resulting patterns again transformed through ***T***_1_ to produce the patterns in ROI Y.

Figure 2b shows the second scenario where patterns are associated through linear multidimensional relationships, with vertices in ROI X being uncorrelated to each other but transformed to ROI Y through a matrix ***T***, so that ***Y*** = ***XT*** + ***E***, where ***T*** is of size *n*_*x*_ × *n*_*y*_. Scenario 3 in Figure 2c illustrates patterns associated through nonlinear multidimensional relationships, with patterns in ROI X first being transformed through ***T***_**0**_, as a linear mapping, then the resulting output entries passed through a nonlinear function, in this case sigmoid or tangent hyperbolic functions (these activation functions are widely used in NNs and may better reflect how neurons summarise information than unbounded activation functions (Sharma et al., 2017)). The resulting patterns are then transformed through ***T***_**1**_ to produce patterns in ROI Y, so that 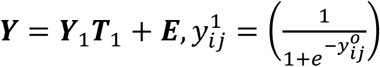, or 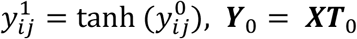, where E is a zero-mean Gaussian noise of size *n*_*t*_ × *n*_*y*_. To avoid bias due to the particular nonlinear function employed in our simulations, we used two nonlinear functions to create the nonlinearity between the patterns (tanh, which is also used in the ANN, and a different function, sigmoid).

#### 2.2.1 Simulation Parameters

To assess the performance of nTL-MDPC, we varied the critical parameters, namely the number of trials, vertices, and signal-to-noise ratios (SNR), within realistic ranges. The numbers of vertices used were either 5 or 15, being matched with the minimum and maximum number of vertices obtained from the implementation of our clustering approach on our EEG/MEG dataset (Rahimi et al., 2022b). To report the final connectivity values, the average of 100 simulations was calculated for every set of parameters.

#### 2.2.2 Scenario 1: Checking for spurious connectivity between two independent patterns

To contrast TL-MDPC and nTL-MDPC on their likelihood to detect false positives, we computed the connectivity between patterns with no relationship (as in Figure 2a). To do so, two pattern matrices with random noise were generated independently using normal distributions (mean=0 and std=1) across 30, 50, 100, 150, and 300 trials.

#### 2.2.3 Scenario 2: Testing the methods’ ability to capture linear multidimensional dependency between two patterns

We then compared TL-MDPC and nTL-MDPC on their ability to detect multidimensional linear relationships, as shown in Figure 2b. To do so, we produced patterns ***X*** through a normal distribution (mean=0, and std=1). For the transformation matrix, we generated matrices using a normal distribution (mean=0, and std=1) with different degrees of sparsity (varying from 10% of the matrix size to 100%, with 10% as the step size). ***Y*** was computed through multiplying ***X*** and ***T***. Different levels of noise were then introduced from a zero-mean normal distribution with a varying standard deviation (std=10^std_pow^, where std_pow ∈[-2, -1.5, -1, -0.5, 0, 0.5, 1, 1.5, 2]). We used 50, 150, and 300 as the number of trials.

#### 2.2.4 Scenario 3: Testing the ability of each method to capture nonlinear multidimensional dependency between two patterns

Constructing nonlinear relationships to compare the methods is not as straightforward as making the patterns in the two previous scenarios, as there are infinitely many ways to simulate a nonlinear function. Here, we chose a method that is easily tractable yet inspired by our knowledge of neuronal interactions. The creation of ANNs was inspired by how neurons function in the brain and how they learn through neural plasticity (Hebb, 2005; McCulloch and Pitts, 1943). In this framework, a unit or node reflects a neuron (or populations of neurons), the synapses are represented as weighted connections between these nodes, and the firing of a neuron (or population of neurons) is determined by the weighted input from other neurons passed through a nonlinear transfer function (Hebb, 2005; McCulloch and Pitts, 1943). Thus, we mimicked this process to create activation patterns with a nonlinear relationship that could reflect the kinds of relationships found in the brain. First, we generated ***X*** and ***T***_0_ using a normal distribution (mean=0, and std=1). The output of the first layer is gained through multiplication of ***X*** with ***T***_0_. Second, this (linear) output is then fed through a nonlinear sigmoid or tangent hyperbolic function. Third, the resulting matrix again passes through another transformation matrix, ***T***_1_, yielding ***Y***. We introduced zero-mean noise from a normal distribution with a variable standard deviation (std=10^std_pow^, where std_pow ∈[-2, -1.5, -1, -0.5, 0, 0.5, 1, 1.5, 2])). The number of trials, vertices, and replications are the same as section 2.2.3. As in scenario 2, both *T*_0_ and *T*_1_ were generated with different degrees of sparsity.

### 2.3 Comparing nTL-MDPC to TL-MDPC in a real EEG/MEG dataset

We applied nTL-MDPC to an existing dataset (for more detail see S.-R. Farahibozorg, 2018; Rahimi et al., 2022a) to investigate the task modulation of semantic brain networks in visual word recognition, contrasting a semantic decision (SD) task requiring deep semantic information retrieval with a lexical decision (LD) task that only requires shallow semantic processing. We assessed whether nTL-MDPC captures different or more functional connections in the semantic network, as compared to TL-MDPC and coherence.

#### 2.3.1 Participants

Our EEG/MEG dataset contains recordings from 18 healthy native English speakers (mean age 27.00±5.13, 12 female) with normal or corrected-to-normal vision. We used the same dataset as in our prior study to compare the outcome of the new method to the linear version of TL-MDPC (Rahimi et al., 2022a). The experiment was approved by the Cambridge Psychology Research Ethics Committee and volunteers were paid for their time and effort.

#### 2.3.2 Stimuli and procedure

The experiment included 250 words and 250 pseudowords. It comprised four blocks presented in a random sequence. One of the four blocks used a lexical decision (LD) task and the other three a semantic decision (SD) task. In the LD block, participants were asked to decide if the presented stimulus was referring to a word or pseudoword. In SD blocks, they were required to decide whether the presented word was referring to a certain category of words, namely “non-citrus fruits”, “something edible with a distinctive odour” and “food containing milk, flour or egg”. 10% of stimuli belonged to these target categories and required a button press response. As in our previous studies, we only analysed brain responses to real, non-target words. Each stimulus was presented for 150ms, with an average SOA of 2400ms.

#### 2.3.3 Data Acquisition and Pre-processing

As we used the same dataset from (S.-R. Farahibozorg, 2018; Rahimi et al., 2022b, 2022a) and intended to directly compare the results of the two methods, data acquisition and pre-processing steps are exactly the same. MEG/EEG recordings were collected simultaneously using a Neuromag Vectorview system (Elekta AB, Stockholm, Sweden) and MEG-compatible EEG cap (EasyCap GmbH, Herrsching, Germany) at the MRC Cognition and Brain Sciences Unit, University of Cambridge, UK. MEG was acquired via a 306-channel system consisting of 204 planar gradiometers and 102 magnetometers. EEG was collected via a 70-electrode system with an extended 10-10% electrode layout. Data sampling rate was 1000Hz.

We used MEGIN Maxwell-Filter software to apply signal source separation (SSS) with its spatiotemporal extension to remove noise from spatially distant sources and to compensate for small head movements (Taulu and Kajola, 2005). We used the MNE-Python software package (Gramfort et al., 2014, 2013) to perform the preprocessing and source reconstruction. The raw data from each participant was visually checked and bad EEG channels were selected for interpolation (max=9 channels per person, min=0, mean=2.85). A finite-impulse-response (FIR) bandpass filter between 0.1 and 45 Hz was applied, as well as FastICA algorithm to remove eye movement artefacts (Hyvarinen, 1999; Hyvärinen and Oja, 2000). Afterwards, epochs were created using the data from 300ms pre-stimulus to 600ms post-stimulus.

#### 2.3.4 Source Estimation

L2-Minimum Norm Estimation (MNE) (Hämäläinen and Ilmoniemi, 1994; Hauk, 2004) was used for source estimation. We assembled inverse operators based on a 3-layer Boundary Element Model (BEM) of the head geometry, gained from structural MRI images. For this purpose, we assumed sources to be perpendicular to the cortical surface (“fixed” orientation constraint). We obtained the noise covariance matrices using 300ms-baseline periods and then selected the best choice from a group of methods included in MNE-Python (‘shrunk’, ‘diagonal_fixed’, ‘empirical’, ‘factor_analysis’) (Engemann and Gramfort, 2015). MNE-Python’s default SNR = 3.0 was used to regularise the inverse operator for evoked responses. The source signals from each participant were then morphed to the standard Freesurfer brain (fsaverage).

#### 2.3.5 Regions of Interest

We used six regions of interest including left and right ATL, left IFG, left PTC, left AG and left PVA to study connectivity within the semantic network. These regions were identical to our prior studies and were constructed using the Human Connectome Project (HCP) parcellation (Glasser et al., 2016). For more details refer to our previous studies (Rahimi et al., 2022a, 2022b).

#### 2.3.6 Leakage

Leakage is an issue for EEG/MEG source signals, leading to limited spatial resolution (Hauk et al., 2022). As introduced in Rahimi et al. (2022a), here we present leakage indices to explain leakage among our six regions. These leakage indices are identical to those presented in Rahimi et al.(2022b), and are simply displayed on a brain diagram here to help interpretation of the current results. Leakage between brain regions can be described by a linear vertex-to-vertex transformation, given by the corresponding sub-matrix of the resolution matrix (Hauk et al., 2022). Figure 3a shows the different steps taken to obtain the leakage index. First, the resolution matrix is computed and then the relevant point spread functions (PSFs) (Hauk et al., 2011; Liu et al., 2002) are extracted. Second, non-homogenous activation patterns are generated for each ROI, to be used at each vertex to weigh their corresponding PSFs. The weighted sum of the PSFs are then computed. Third, the leakage is summarised by taking the absolute values and summed across vertices per ROI. Fourth, the leakage index is calculated by dividing the summed leakage of an ROI to itself by the summed leakage from another ROI. This stage was replicated 100 times for each PSF and participant, and the results were averaged to create the final leakage indices. Figure 3b represents our leakage matrix. The *i*th column shows how much all other ROIs leak into the *i*th ROI relative to the *i*th ROI’s leakage into itself. To have a better understanding of the matrix, we considered leakage values between 0-0.2/0.2–0.4/0.4–0.6/0.6–0.8/0.8–1 to reflect low/low-medium/medium/medium-high/high leakage.

**Figure 3.**
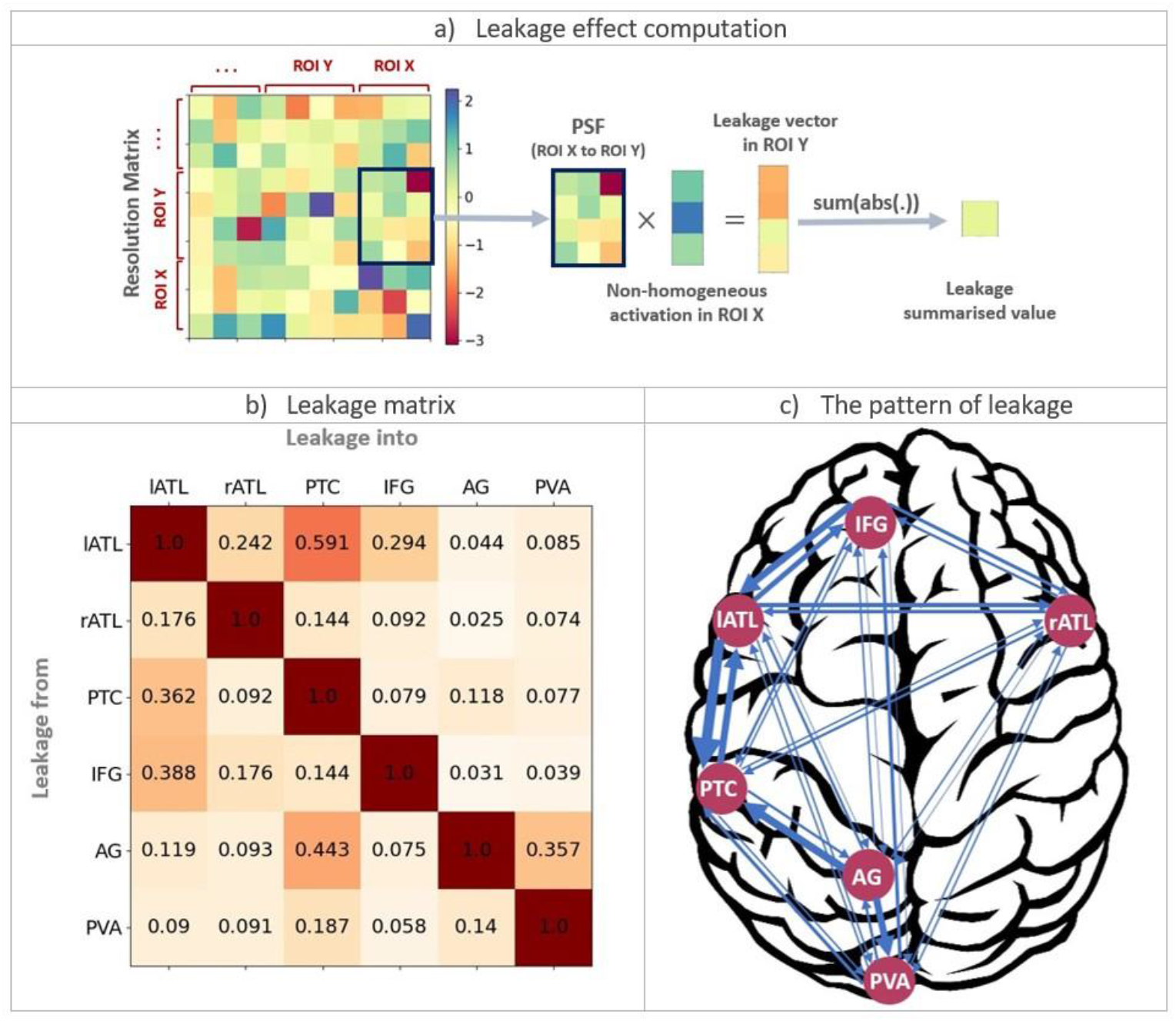
Leakage indices for multidimensional activation patterns. a) Schematic description of how to compute the non-homogeneous leakage from ROI X into ROI Y using the resolution matrix and point-spread-functions (PSFs) as well as non-homogeneous activation vectors. b) The leakage matrix for our six ROIs. c) The pattern of leakage across the semantic network. The width of the arrows reflects the leakage indices in b)(obtained from Rahimi et al., 2022b).

The leakage matrix indicated that all region pairings except for lATL to PTC, have medium or lower leakage. The highest leakage is between two nearby areas, lATL and PTC. Note that high leakage is likely for any study investigating connectivity between these regions, and that we combined EEG and MEG to minimise the leakage. Figure 3c represents the strength of the directed leakage indices between all pairs, reflected in the width of the arrows. We will compare the networks obtained from our real data analysis with the pattern of leakage across the semantic network to consider the possible confounding effect of leakage on our results.

#### 2.3.7 Application of nTL-MDPC to real EEG/MEG data

We performed nTL-MDPC to predict ***X*** from ***Y*** and ***Y*** from ***X*** at every 25ms from 100ms pre-stimulus to 500ms post-stimulus, using an ANN with a tangent hyperbolic activation function, and reported the EVs (averaged across the two directions) as the final metric. EVs for every pair of ROIs and across latencies are presented in TTMs (Rahimi et al., 2022b). In any TTM, each row indicates statistical relationships between ***Y*** at a specific time point and *X* over the whole time period, while columns show statistical relationships between ***X*** at a certain time point and ***Y*** across the whole time period, enabling us to display connectivity at different time lags.

To avoid any potential bias due to the different number of trials between SD and LD tasks (Bastos and Schoffelen, 2016), the TTMs of the three SD blocks were computed independently and then averaged for a better comparison with LD. SD and LD connectivity were then contrasted using cluster-based permutation tests, implemented in MNE python (Maris and Oostenveld, 2007). We implemented two-sided t-tests with the alpha-level of 0.05 and 5000 randomised repetitions. To avoid spurious or small clusters, we only presented clusters whose size were greater than 2% of the TTMs total size, i.e., with more than 12 elements.

## 3 Results

### 3.1 Simulation results

#### 3.1.1 Scenario 1: Independent patterns

demonstrates connectivity scores (EV, y-axis) for the case where there is no true relationship between two patterns (as in Figure 2a), with the x-axis showing different numbers of trials and different curves representing different numbers of vertices for both panels. Figure 4a shows the connectivity results of MDPC, while Figure 4b indicates the nMDPC outcomes. For both methods, all values are close to zero, suggesting that neither method is prone to false positive errors when no relationship exists.

**Figure 4.**
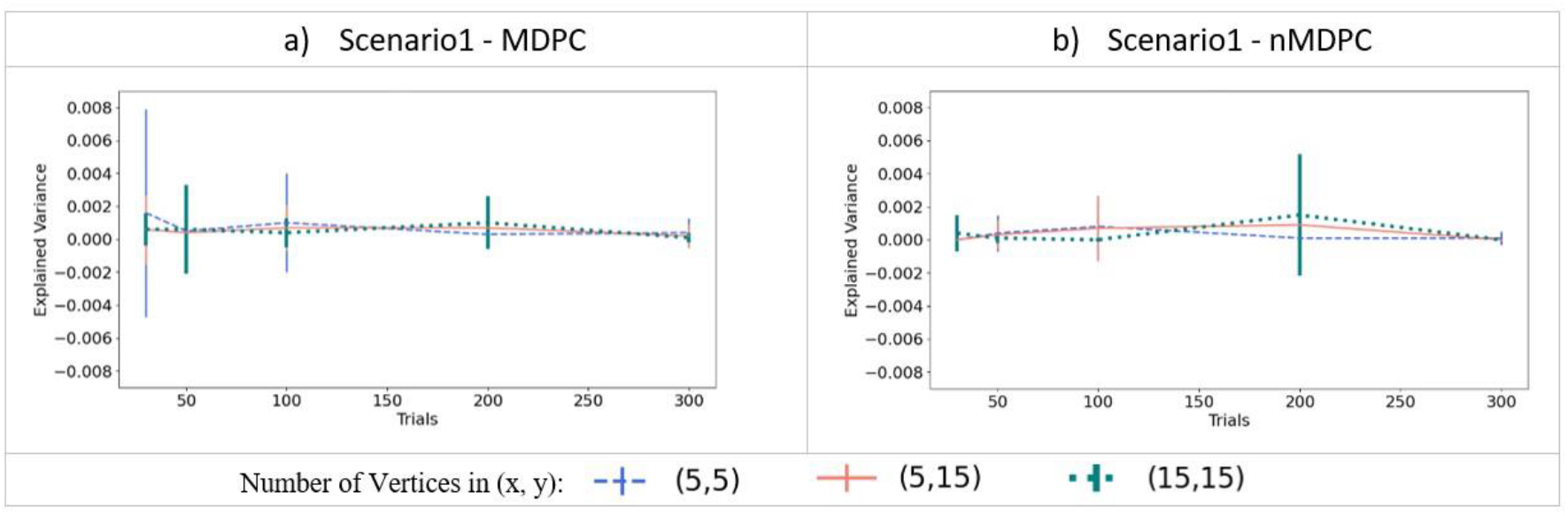
Connectivity values (explained variance, y-axis) for two independent patterns. a) Connectivity for (linear) MDPC as a function of different numbers of trials, with different curves representing different combinations of numbers of vertices in ROI X and ROI Y. b) Similar to a), but for nMDPC. The values are close to zero indicating that neither MDPC nor nMDPC methods are prone to false positive errors for random patterns. Note that while all EVs and their means were positive (as negative EVs were replaced by zeros), error bars based on standard deviations can still extend below zero.

#### 3.1.2 Scenario 2: Linear multidimensional dependency between two patterns

We then tested how well the MDPC and nMDPC methods perform in the case where multidimensional patterns are only linearly related. Figure 5 demonstrates the connectivity (EV, y-axis) identified in this scenario, with the x-axis representing different SNRs, different curves indicating different trials, and three panels showing different numbers of vertices. Red curves show the result of the MDPC method and blue curves represent the nMDPC results. For both methods and in all cases, EV reaches values between 0.8 to 1 for SNRs above 25db and is close to zero for SNRs smaller than -10db. For intermediate SNRs the two methods perform similarly, with EV rising for increasing numbers of trials. Thus, both linear and nonlinear MDPC similarly capture true linear relationships between patterns to a similar degree, with MDPC producing slightly greater EV values.

**Figure 5.**
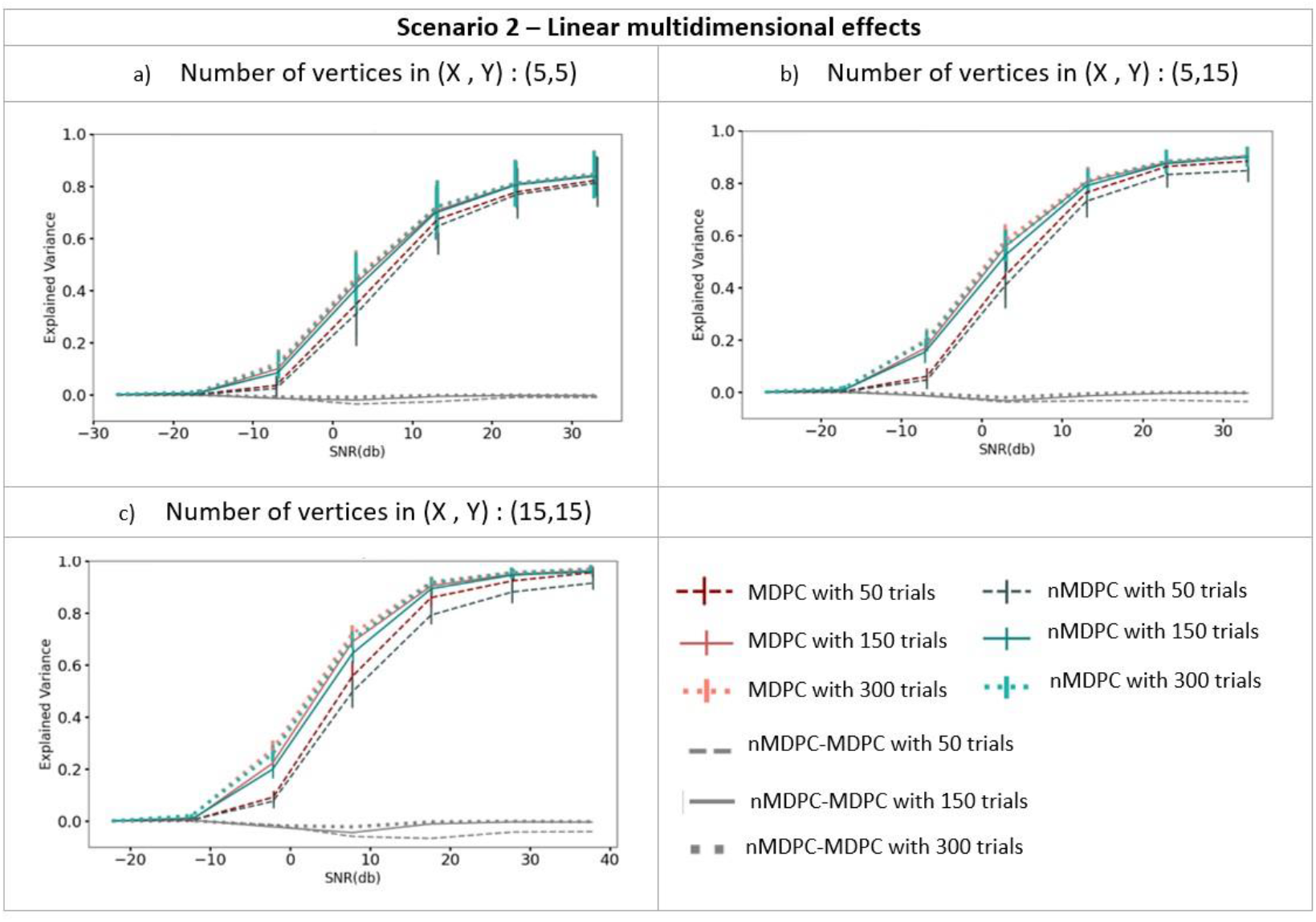
Connectivity metrics for linear multidimensional effects assessed using MDPC and nMDPC. All panels show connectivity scores (explained variance, y-axis) between patterns that have a true linear multidimensional dependency, for different SNRs (x-axis), three different numbers of trials (MDPC approach: red curves, nMDPC approach: blue curves) and different number of vertices (three panels). All cases show that EV (explained variance) reaches value above 0.8 for SNRs above 25db, and is close to zero for SNRs smaller than -10db. In all panels, both linear and nonlinear MDPC show similar results, with MDPC producing greater EV.

#### 3.1.3 Scenario 3: Nonlinear multidimensional dependency between two patterns

Figure 6 shows the connectivity (EV, y-axis) between pairs of multidimensional patterns with a nonlinear relationship (see section 2.2.4), for different SNRs (x-axis), different trials (indicated by line type), and different numbers of vertices (three panels). We used two different functions to generate nonlinear dependencies, sigmoid (shown in Figure 6a, c, and e) and tangent hyperbolic (shown in Figure 6Figure 6b, d, and f). EV is zero for all cases at very low SNRs (<-10dB). Unlike for linear dependencies above, the EV does not approach 1 at high SNRs. For a given number of trials, nonlinear MDPC generally outperforms its linear counterpart. However, the difference between methods is not large and is around 0.1. Figure 6e and 5f show that the worst performance for both methods is obtained for the largest number of vertices and the lowest number of trials. Interestingly, in this case the linear method outperforms the nonlinear one. This is likely due to the known requirement of ANNs for a lot of training data (Hagan and Demuth, 1999). Thus, while linear MDPC provides a good approximation to the nonlinear multidimensional relationships when a large number of parameters needs to be estimated from a small number of trials, in general, nMDPC captures more variance. This improved ability of nMDPC to detect additional nonlinear relationships, could allow the identification of additional functional connections in brain data, including connectivity between more regions or across additional time lags.

**Figure 6.**
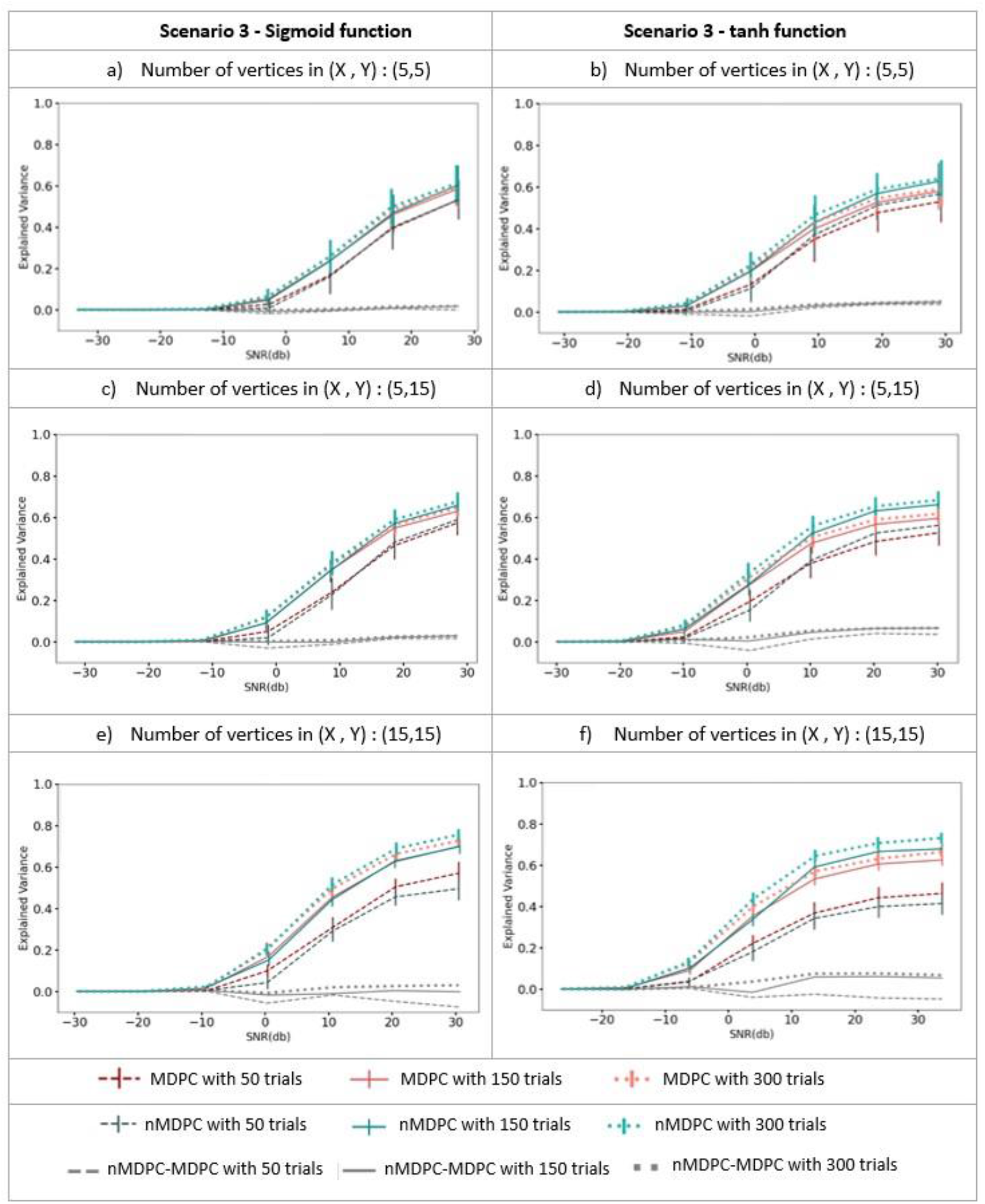
Connectivity metrics for nonlinear multidimensional effects using MDPC and nMDPC. a, c, e) Connectivity scores (explained variance, y-axis) between patterns with nonlinear multidimensional dependency, for different SNRs (x-axis), three different numbers of trials (MDPC approach: red curves, nMDPC approach: green curves) and different number of vertices (three panels). b, d, f) Same as a, c, e) but with a tangent hyperbolic activation function. Linear MDPC provides a good approximation to these nonlinear scenarios in most cases, and even performs slightly better than nMDPC for low numbers of trials and large numbers of vertices. However, nMDPC generally explains more variance particularly for larger numbers of trials.

### 3.2 Comparison of nTL-MDPC and TL-MDPC in a real EEG/MEG dataset

To test whether nTL-MDPC can confirm and extend our previous neural functional connectivity results, we applied nTL-MDPC to data from an EEG/MEG experiment contrasting a semantic decision and lexical decision task (R. Farahibozorg, 2018; Rahimi et al., 2022a). We compared the results of nTL-MDPC to TL-MDPC analyses of this dataset, previously presented in Rahimi et al.(2022b). Specifically, we asked whether and how the connectivity of semantic network is modulated by task demands, and if nTL-MDPC detects more or different dynamic connectivity compared to TL-MDPC.

Figure 7 illustrates the interpretation of nTL-MDPC and TL-MDPC results, in the form of TTMs for one pair of ROIs, in this case PTC and IFG, two regions putatively involved in semantic control (Rahimi et al., 2022b). Figure 7a and 7b show results for TL-MDPC and nTL-MDPC, respectively, separately for the semantic decision task (SD) and lexical decision (LD) task, as well as their statistical comparison (using cluster-based permutation tests). An element of the TTM at coordinate (*x,y*) describes the relationship between patterns in IFG at latency *x* and PTC at latency *y*. Note that these statistical relationships do not imply a directionality of the corresponding connectivity, even at non-zero time lags. As expected, we found greater connectivity with both approaches for the more semantically demanding SD task compared to the LD task. The separate TTMs for SD and LD have a similar shape, with the largest values concentrated around the diagonal. However, the statistical comparison between the tasks highlights the more reliable identification of connectivity for nTL-MDPC along the diagonal, as well as at larger time lags, particularly at later latencies. Interestingly, the area of significant differences appears to “fan out” over time, i.e., connectivity persists over longer time lags at later latencies in both upper and lower diagonal parts, possibly reflecting bidirectional or recurrent information flow (Clarke et al., 2015, 2011; Kietzmann et al., 2019; McClelland and Rumelhart, 1989; Rogers et al., 2021; Rogers and McClelland, 2014). In this example, the nonlinear method also showed modulation of connectivity during the baseline interval, which is confined to near the diagonal (i.e., close to zero-lag). Effects during the baseline were previously identified between some ROI pairs using the linear method. These may reflect a modulation of baseline activity due to differences in expectation and preparation when using tasks with different demands in a blocked experimental design.

**Figure 7.**
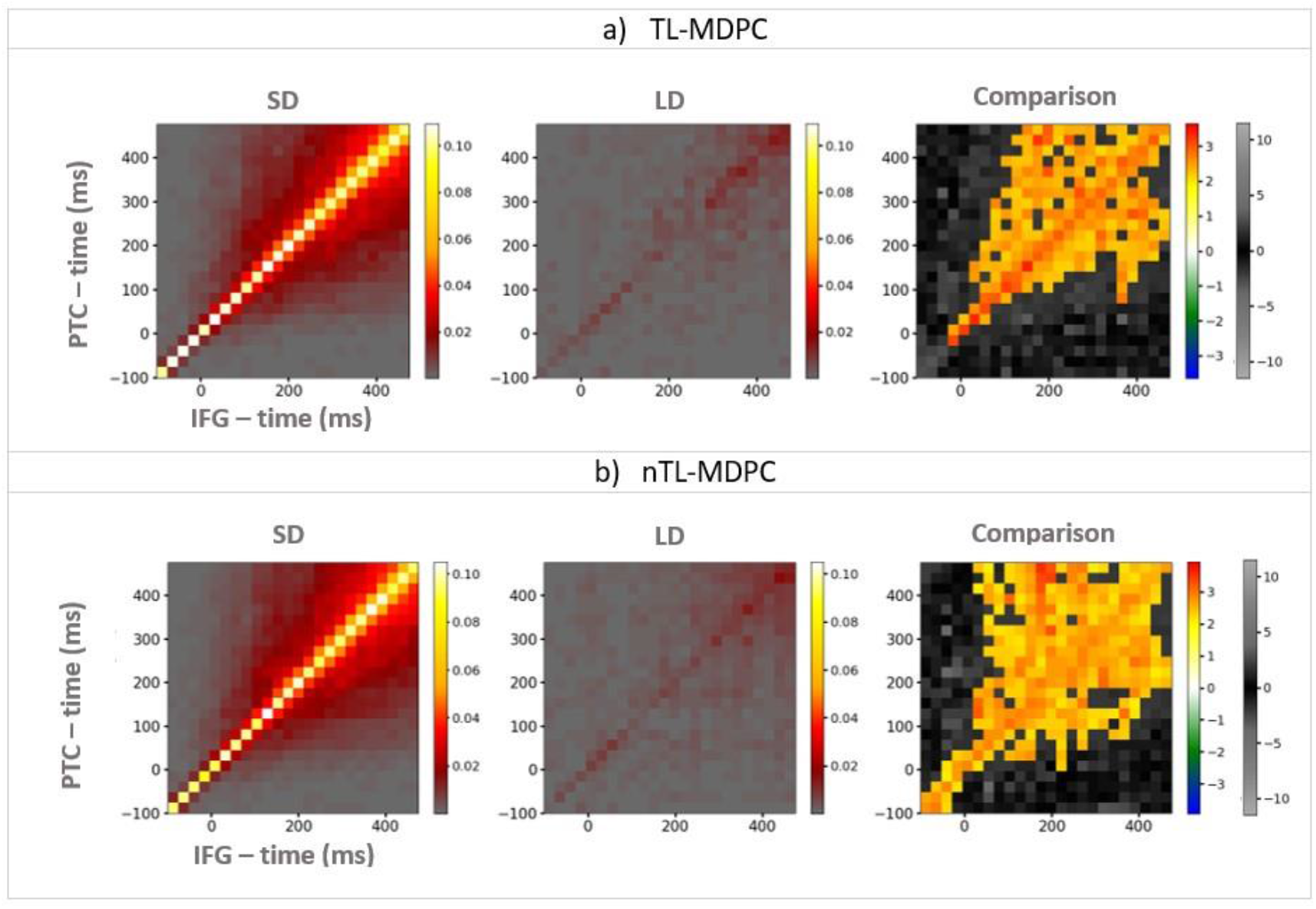
An example of TTMs showing the connectivity between PTC (y-axis) and IFG (x-axis), for the semantic decision (SD) task (the left column), the lexical decision (LD) task (the middle column), and their comparison (i.e., connectivity that is greater when there are greater semantic demands; right column). a) TTMs for TL-MDPC, b) TTMs for nTL-MDPC. Colorbars show connectivity scores (explained variance) for the first two left columns. For the third column, the hot and cold color bar highlights significant effects obtained from the cluster-based permutation test, whereas the gray-scale color bar shows non-significant t-values (this color bar is the same across all Figures). With both methods, the greatest connectivity accurs around the diagonal, however, the statistical comparison between the tasks reveals more reliable modulation of connectivity using nTL-MDPC along the diagonal as well as at larger time lags, particularly at later latencies.

#### 3.2.1 Detecting connectivity within the semantic network across time with nTL-MDPC

We computed the functional connectivity of all possible pairs of the six semantic ROIs using nTL-MDPC and TL-MDPC. Figure 8 represents the inter-regional connectivity matrix (ICM) (Rahimi et al., 2022b), a summary of the semantic decision versus lexical decision TTMs for all ROI comparisons, with the upper diagonal (green area) showing the nTL-MDPC results, and the lower diagonal (blue area) displaying the TL-MDPC results. Generally, both methods found greater connectivity in the SD compared to the LD task for the same pairs of regions. Specifically, eleven pairs were modulated by task differences: lATL-rATL, lATL-PTC, lATL-IFG, rATL-PTC, rATL-IFG, rATL-AG, rATL-PVA, PTC-IFG, PTC-AG, PTC-PVA, and AG-PVA. The connectivity between core semantic regions including lATL-rATL, lATL-PTC, lATL-IFG, rATL-PTC, PTC-IFG, as well as AG-PVA, fans out from the diagonal from early time points. Connectivity for rATL-IFG was mostly confined around the diagonal, and rATL-AG along with rATL-PVA revealed late effects. However, nTL-MDPC captured more connectivity in some cases. For instance, for some pairs it revealed more modulations across more time lags, e.g., between lATL-rATL, lATL-PTC, rATL-PTC, PTC-IFG. However, these differences were subtle and when the two methods were contrasted directly, there were no significant differences.

**Figure 8.**
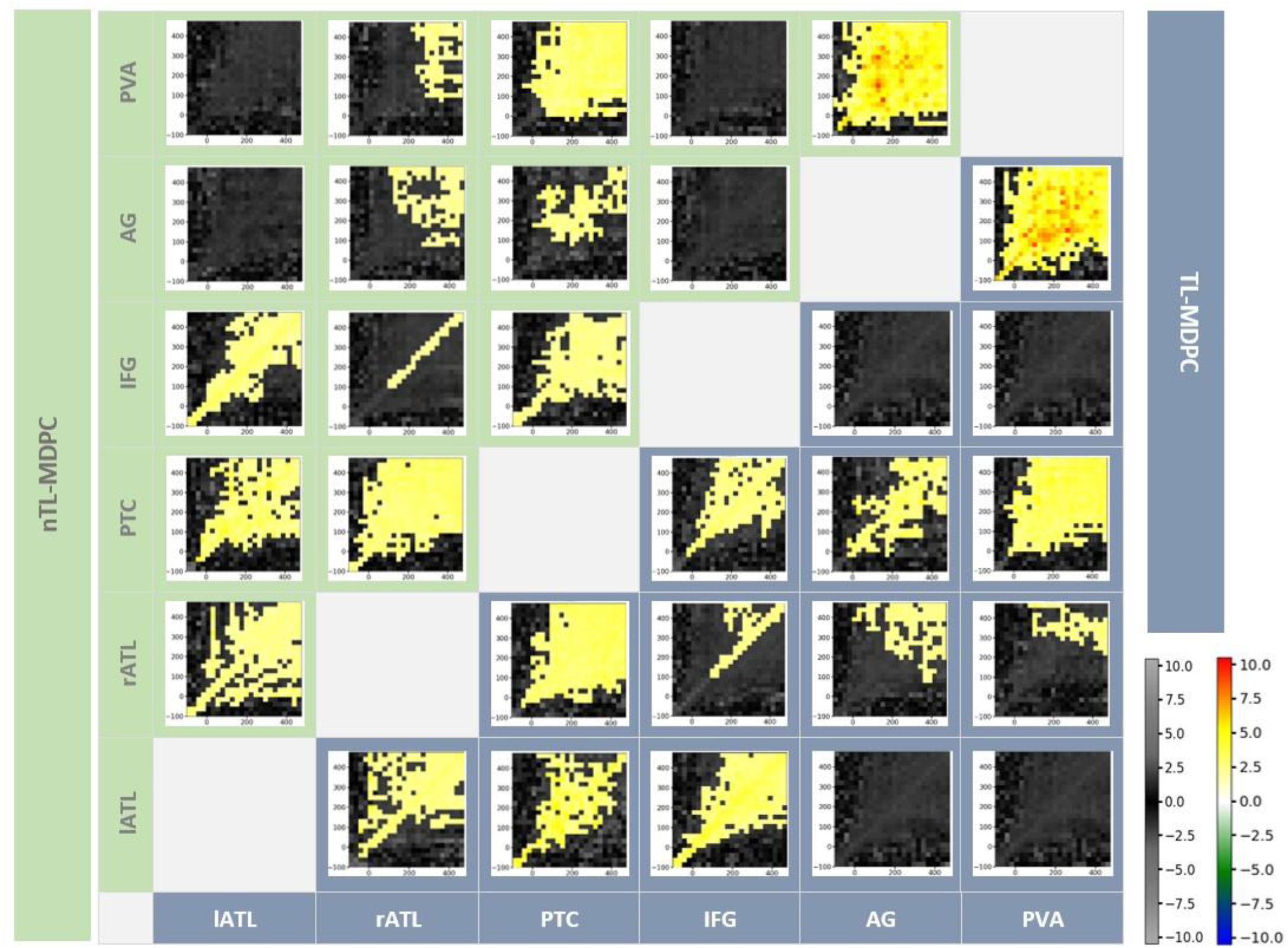
Inter-regional Connectivity Matrix (ICM) for semantic network modulations in the brain – the upper diagonal (grey shaded area) shows nTL-MDPC TTMs and the and lower diagonal (blue shaded area) shows TL-MDPC TTMs. Each TTM reflects EVs for a pair of region, ROI X and ROI Y, and across times. All modulations with both methods showed greater connectivity for SD than LD. Cluster size was thresholded at 2% of TTMs size (24*24). Using both methods, we found rich modulations between semantic control and representation regions. The hot and cold color bar highlights significant effects obtained from the cluster-based permutation test, whereas the gray-scale color bar shows non-significant t-values. The color bars are the same across all Figures.

As multiple different methods have now been used to assess connectivity within the same dataset, we summarised the networks extracted using these different methods in Figure 9. This demonstrates the results from: 1) coherence (Rahimi et al., 2022a), 2) TL-UDC (Rahimi et al., 2022b), 3) TL-MDPC (Rahimi et al., 2022b), and 4) nTL-MDPC analyses, for early (50-250ms) and late (250-500ms) time windows. TL-UDC is the unidimensional counterpart of TL-MDPC (Rahimi et al., 2022b), in which all the time courses of the vertices within each ROI are collapsed into one. The ccoherence (Rahimi et al., 2022a) between each pair of ROIs was computed for each time window in four frequency bands, namely theta (4-8Hz), alpha (8-16Hz), beta (16-26Hz), and gamma (26-36Hz).

**Figure 9.**
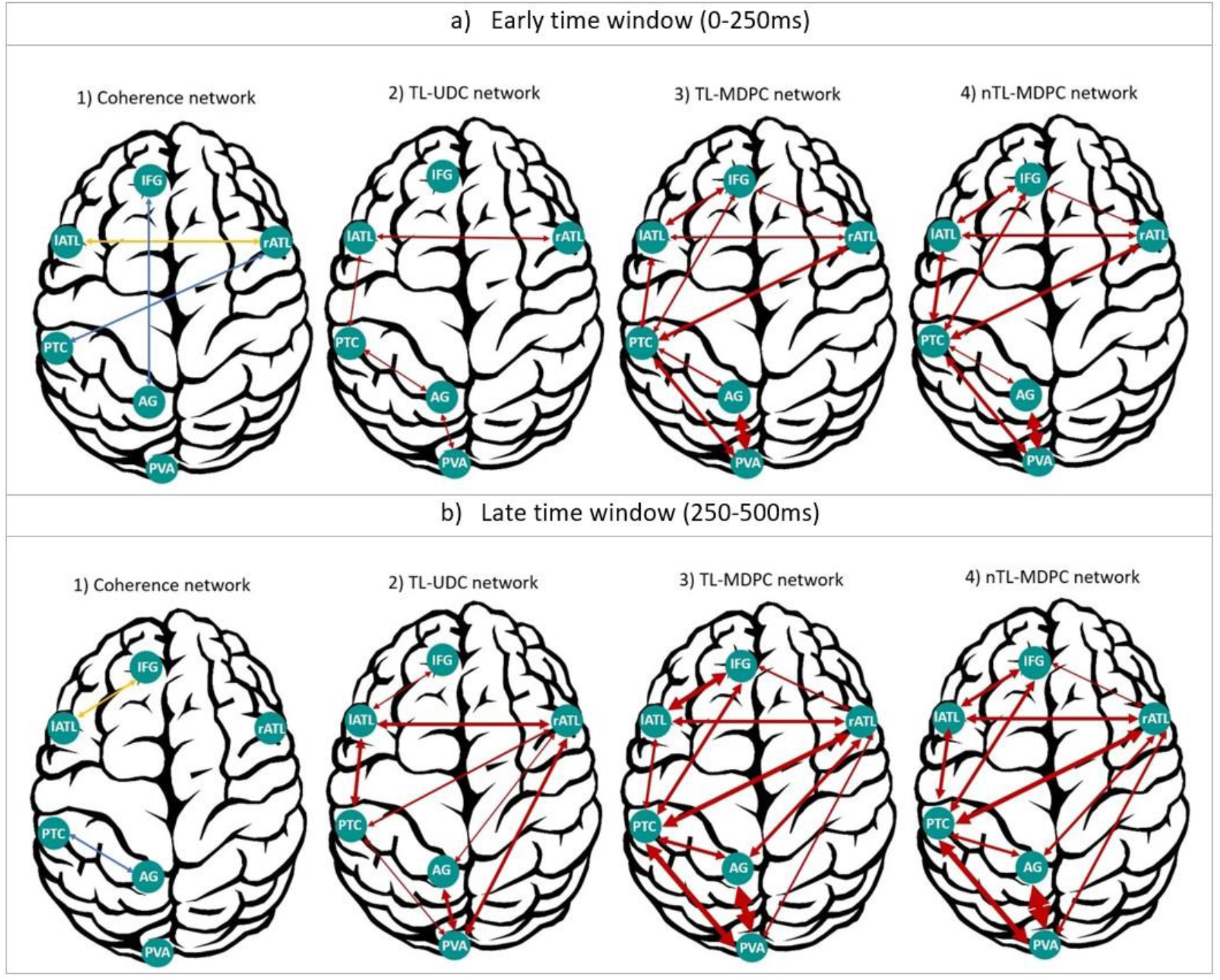
Summary of our cortical functional connectivity analyses of EEG/MEG data using four different connectivity approaches. a) Networks extracted for an early time window (0-250ms), through 1) coherence (Rahimi et al., 2022a), 2) the UDC method (Rahimi et al., 2022b), 3) MDPC (Rahimi et al., 2022b), and 4) nMDPC. b) Same as a), but for a late time window (250-500ms). For coherence, the blue connections represent dependencies specific to the gamma band, and yellow ones show the connections consistant across the alpha, beta and gamma bands. For the other three methods, arrows represent the summed significant t-values in each time window as a metric of connection intensity, with thicker arrows reflecting more intense connections (higher summed t-values) and vice versa. Overall, TL-MDPC reveals more connections compared to the two unidimensional methods, but TL-MDPC and nTL-MDPC produced the same pattern of connectivity with only a few differences. Despite this similarity in the areas found to be connected, nTLMDPC finds connections more reliably across more time lags. Using nTL-MDPC, we found a connection intensity in a higher band for lATL-rATL and PTC-IFG connections at the early time points, and for lATL-PTC, PTC-IFG, and rATL-PVA at the later time points. However, lATL-IFG and PTC-AG had a connection intensity in a higher band in the later time window for the linear TL-MDPC method.

In this figure, we represent the connections specific to the gamma band in blue (i.e., rATL-PTC, IFG-AG at the first time window, and PTC-AG at the second time window), and connections consistent across the three frequency bands (i.e. alpha, beta, and gamma) in yellow (i.e., lATL-rATL, at the first time window, and lATL-IFG at the later time window). To summarise the ICM results across time lags for TL-UDC, TL-MDPC, and nTL-MDPC, we summed the significant t-values in each time window as a metric of connection intensity. This reflects the strength of the connections found and the identification of significant connectivity across longer time lags, indicating more reliable identification of connections with higher values, as well as greater reliability across participants. We then represented these values using arrows with different widths (with the weakest connectivity reflected by the thinnest lines, as in rATL-IFG using nTL-MDPC at the earlier time window, and the strongest connectivity represented by the thickest line, as in AG-PVA using nTL-MDPC at the later time window).

In general, the two unidimensional connectivity methods revealed fewer and weaker task modulations of connectivity, while both multidimensional methods detected richer connectivity across the whole semantic network across the whole latency range. Both multidimensional methods can identify rich interactions between the four core semantic representation and control regions (lATL, rATL, PTC, and IFG) starting at early time points (around the onset of the stimulus) and becoming stronger across the time window. Even with methods designed to identify nonlinear relationships, the connectivity of AG is relatively sparse. It only shows connectivity with PVA and PTC from earlier time points, and with rATL at later latencies. The general pattern of results was similar for TL-MDPC and nTL-MDPC, with both distinguishing a core semantic network comprising lATL, rATL, PTC and IFG from a posterior visual attention network consisting of AG, PVA and PTC, with overlap of these two sub-networks in PTC.

While the contrast between our linear and nonlinear results did not yield any significant effects, visual inspection of Figure 9 reveals some subtle but noteworthy differences between the two approaches. For instance, nTL-MDPC finds more connections across more time lags. Using nTL-MDPC, we found slightly stronger connectivity for lATL-rATL and PTC-IFG at early time points, and for lATL-PTC, PTC-IFG, and rATL-PVA at the later time points. However, connectivity was slightly stronger for lATL-IFG and PTC-AG when using the linear TL-MDPC method at late time window. The finding of a greater number of numerically stronger connections for the nonlinear method could suggest some subtle benefits for this method in the overall reliability in the evidence of connectivity between the regions of the semantic network for the two methods. However, this did not result in a significant difference between these methods in our statistical analysis.

## 4 Discussion

We introduced a novel multidimensional functional connectivity method, nTL-MDPC, to capture the nonlinear dependencies between event-related EEG/MEG activation patterns across different brain regions and at different time lags. Our simulations revealed that nTL-MDPC is not prone to false positive errors for independent and uncorrelated patterns and is able to capture true linear multidimensional effects between patterns, although in this case the linear method performed slightly better. For true nonlinear dependencies, it it generally outperforming the linear method except for low numbers of trials and high number of vertices. However, this was not the case in the real data analysis, where the linear and nonlinear methods showed similar performance. The results of our new nonlinear approach were generally in line with our previous findings with TL-MDPC, indicating that linear TL-MDPC may provide a good and efficient approximation to nonlinear multidimensional relationships between brain regions in our real EEG/MEG data.

As with TL-MPDC, nTL-MDPC identified two sub-networks in our data, namely a semantic network comprised of four semantic representation and control regions (left and right ATL, PTC, and IFG), and a visual-attention network consisting of PVA and AG. These sub-networks are connected through PTC. These results are in concordance with the controlled semantic cognition (CSC) framework, which emphasises the need for a bilateral semantic hub (ATLs) as well as the interaction between representation and control regions for the task-relevant processing of conceptual information (Lambon Ralph et al., 2016). Across our analyses, the ATLs show early activation and rich connectivity within the semantic network that are modulated by semantic task demands. In contrast, another putative hub region, AG, only showed relative sparse connectivity with posterior brain areas. This supports the findings of Farahibozorg et al. (2022), who used Dynamic Causal Modelling to study word concreteness effects and identified ATL as a connectivity hub in an early (0-250ms) and AG only in a prolonged (0-450ms) time interval. These results suggest a role of AG for example in context integration, episodic memory or attentional processes (Cabeza, 2008; Cabeza et al., 2012; Chambers et al., 2004; Humphreys et al., 2021; Humphreys and Lambon Ralph, 2015; Noonan et al., 2012; Shimamura, 2011; Vilberg and Rugg, 2008; Wagner et al., 2005)

Notably, different methods have different assumptions, and therefore it is useful to look at what is consistent across methods. Across our unidimensional (Rahimi et al., 2022a) and multidimensional (Rahimi et al., 2022b) analyses we revealed early connectivity between left and right ATLs. All analyses showed stronger connectivity between left and right ATLs with stronger semantic demands. Interestingly, both the linear and nonlinear multidimensional methods found rATL connectivity with IFG and PTC in the left hemisphere. This result was not seen with the unidimensional methods, and this cross-hemisphere connectivity is unlikely to be due to leakage. While previous studies have usually observed a left-lateralisation of ATL activation for verbal compared to non-verbal stimuli, right-hemispheric ATL activation is commonly observed in semantic tasks as well (Jackson, 2021; Rice et al., 2015; Visser et al., 2010), in particular in EEG/MEG studies (Dhond et al., 2007; Farahibozorg et al., 2022; Rahimi et al., 2022a). Our results confirm that both left and right ATL are critical for semantic cognition and contribute more in semantically more demanding tasks (Rahimi et al., 2022a; Stefaniak et al., 2022). Some previous studies suggested that rATL might be involved in more semantically demanding tasks (Jung et al., 2019; Stefaniak et al., 2020), which would explain why we observed stronger rATL connectivity with the left hemisphere in the later time window with our multidimensional analyses.

Most of our temporal transformation matrices showed symmetrical patterns with significant effects around the diagonal, which fanned out at later latencies. This suggests that some brain areas might be activated during overlapping time intervals, interacting near-simultaneously over sustained periods of time. Where there is a pattern of sustained symmetrical relationship between regions this may suggest that there is a period of time with a bidirectional information flow between these regions, as shown in prior electrophysiological data (Clarke et al., 2015, 2011; Rogers et al., 2021), possibly reflecting recurrent information flow (Kietzmann et al., 2019; McClelland and Rumelhart, 1989; Rogers et al., 2021; Rogers and McClelland, 2014).

Using both linear (Rahimi et al., 2022b) and nonlinear TL-MDPC, we observed connectivity modulations during the baseline period for some region pairs (e.g., lATL-rATL, PTC-IFG, rATL-PTC). This could be due to the fact that our EEG/MEG data was recorded in a blocked design, affecting brain activity in anticipation of the stimulus and in preparation of a particular type of decision. However, it is important to note that the temporal and spatial extents, and in particular the onsets and offsets, of significant effects from cluster-based permutation tests cannot be determined with precision (Sassenhagen and Draschkow, 2019). The functional significance of these early baseline effects should be investigated in more detail in future studies.

Source leakage is a consequence of the non-uniqueness of the EEG/MEG inverse problem, and therefore affects any processing steps following source estimation, in particular connectivity analysis (Colclough et al., 2015; Farahibozorg et al., 2022; Hauk et al., 2022; Palva et al., 2018). This is a possible confound for all unidimensional (homogenous activation patterns) and multidimensional (non-homogenous patterns) methods. Leakage occurs instantaneously, and therefore methods have been suggested that remove zero-lag dependencies between activation time series to reduce its confounding effect (Colclough et al., 2015; e.g. Nolte et al., 2004). However, if two point sources show true zero-lag connectivity then all vertices that receive leakage from any of the two sources (i.e. where their point-spread functions are not zero) will also show non-zero-lag connectivity, i.e. leakage can still affect non-zero-lag connectivity estimates (Colclough et al., 2015; Farahibozorg et al., 2018; Palva et al., 2018; Rahimi et al., 2022a). In addition, zero-lag connectivity can still be of interest, e.g., for homologous areas in the two brain hemispheres. Both TL-MDPC and nTL-MDPC estimate zero-lag as well as non-zero-lag connectivity (diagonal and off-diagonal elements of the temporal transformation matrix, respectively).

In the present study, we used combined EEG and MEG to gain the optimum spatial resolution and thus minimise leakage (Hauk et al., 2019; Henson et al., 2009; Molins et al., 2008). Furthermore, we represent a quantitative assessment of leakage for our ROIs. While in our previous study we computed leakage indices for the unidimensional case, i.e., homogenous activation patterns, we here extended this approach to the multidimensional case, i.e., non-homogenous activation patterns. The resulting leakage matrix suggests that the highest leakage would be between lATL-PTC, lATL-IFG and PTC-AG. However, the connectivity network shows that the strongest connectivity occurs between 1) AG-PVA, 2) lATL-IFG, rATL-PTC, and PTC-PVA. In particular, we observed reliable connections of rATL with regions in the left hemisphere. Thus, while we cannot rule out that some of our results are affected by leakage as in other similar studies, the overall pattern of our results cannot be explained by leakage alone.

Our approach is a step towards characterising the full dynamic multidimensional information in vertex-to-vertex relationships across brain regions. Both TL-MDPC (Rahimi et al., 2022b) and our new nTL-MDPC applied to dynamic EEG/MEG data allowed more detailed investigation of the semantic network across space and time, compared to previous studies on this network using fMRI (Chiou et al., 2018; Jackson et al., 2016; Jung and Lambon Ralph, 2016) and unidimensional analyses with EEG/MEG (Rahimi et al., 2022a; Sormaz et al., 2017). Our main findings are that 1) in the simulations both methods could identify some linear and nonlinear relationships, with the nonlinear method performing slightly better, and 2) we did not observe significant differences between our methods in our real data analysis. Thus, we cannot report a significant advantage of using a computationally more demanding nonlinear over a simpler linear method, in contrast to the fMRI analyses by Anzellotti et al. (Anzellotti et al., 2017a, 2017b). There are two possible explanations of these results. First, there may be no nonlinear relationships in EEG/MEG data or, second, there may be nonlinear relationships in EEG/MEG data and our linear method may be able to approximate them well enough within the noise level. Future studies should investigate whether this is a general property of EEG/MEG data or whether there are cases for which the nonlinear method is more suitable. As there are many possible ways to perform functional connectivity analyses and we may not be able to take all factors into consideration, we need to know the most effective aspects. For instance, our findings suggest that moving from unidimensional to multidimensional information adds much, yet converting a linear method to a nonlinear version does not. This might help focus our aims on the important aspects for future methods development and applications. It is also possible to decompose nonlinear relationships into linear and nonlinear parts, as for example shown for multivariate connectivity methods (Talebi et al., 2019). This would allow the estimation of linear and nonlinear connectivity components separately within a dataset.

In summary, our novel linear and nonlinear approaches to estimating multidimensional relationships between brain regions have allowed us to reveal novel insights into the dynamics of the semantic brain network, and open new avenues for a richer description of dynamic brain connectivity in the future. We cannot provide any strong evidence that the nonlinear method outperforms the linear one, yet this may not be the case for other datasets. While both methods allowed us to estimate statistical relationships between regions across time lags, they are not directional and therefore do not establish directionality or causality of our observed effects. Furthermore, as for all bivariate methods we cannot determine whether an observed relationship is due to indirect connectivity, e.g., if it is mediated by a third source. In the future, this could be addressed by extending our methods to a multivariate as well multidimensional approach, i.e., considering the full patterns across multiple brain regions simultaneously. This would lead to multivariate autoregressive models which together with the principles of Granger causality could establish directional and even causal relationships between multidimensional time series (Seth, 2010).

## 5 Acknowledgements

This work was supported by intramural funding from the Medical Research Council UK (MC_UU_00005/18), a British Academy Postdoctoral Fellowship awarded to R.L.J. (no. pf170068), and Cambridge University international scholarship awarded to S.R. For the purpose of open access, the author has applied a CC BY public copyright license to any Author Accepted Manuscript version arising from this submission. We also thank Dr Rezvan Farahibozorg for data collection and information on the dataset.

## Supplementary

**Figure S1.**
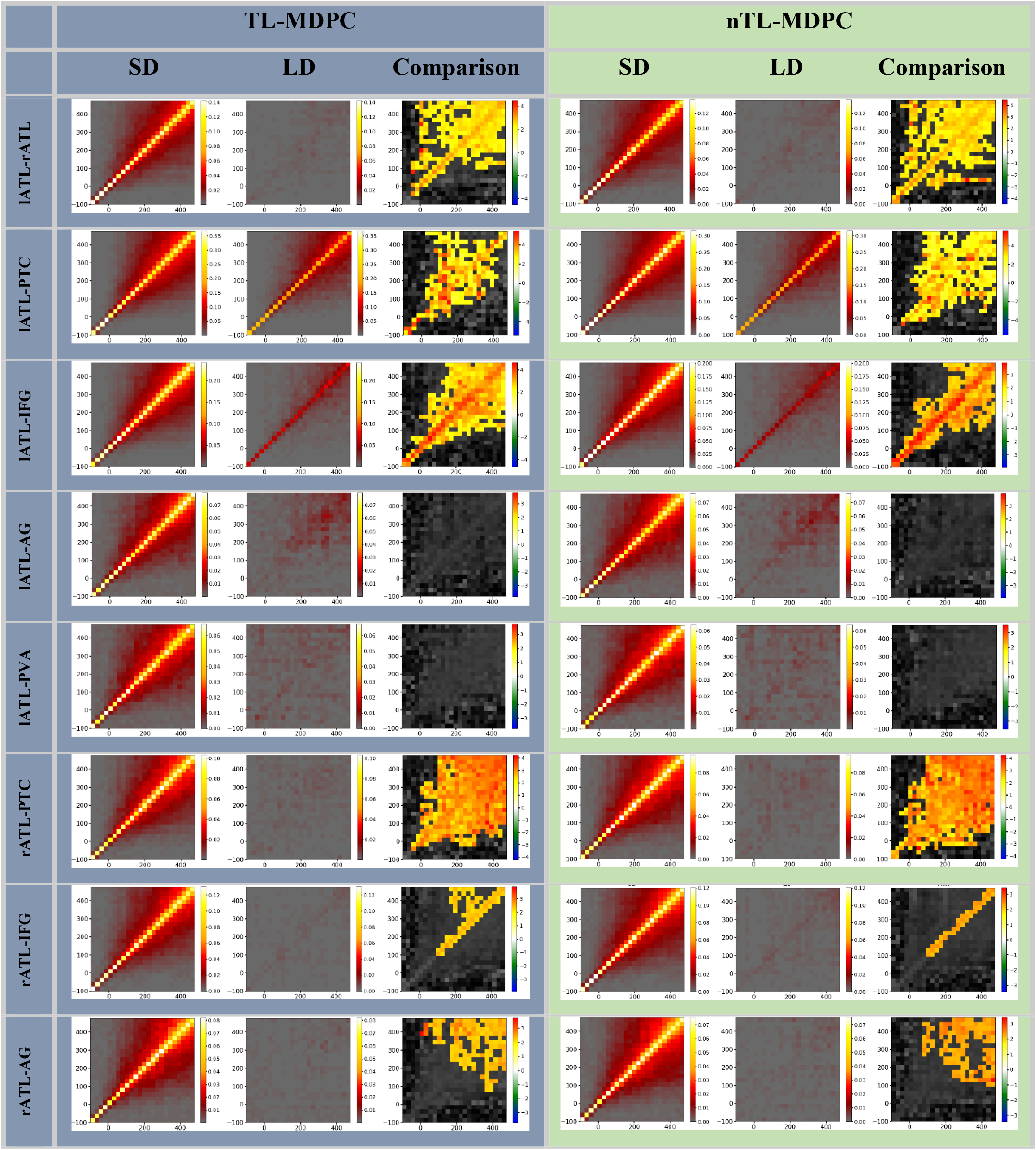
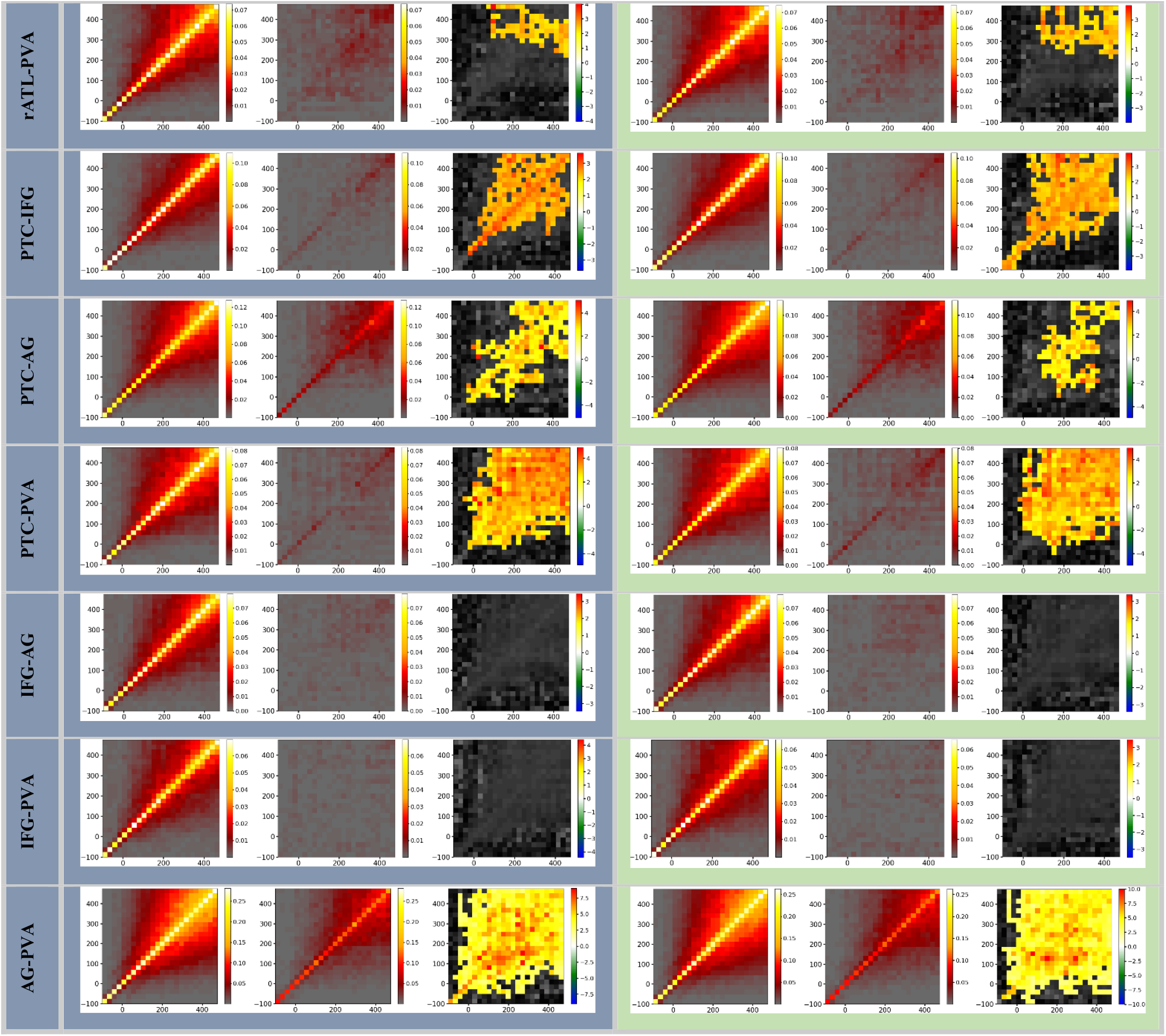
Representation of all TTMs for semantic decision (SD) and lexical decision (LD) tasks, and their comparison, using TL-MDPC (left column) and nTL-MDPC (right column). TTMs for SD and LD are averaged TTMs across participants, and comparison was done using cluster-based permutation test with alpha-level=0.05. All significant contrasts show greater connectivity for SD using MD.

## Footnotes

1 https://scikit-learn.org/stable/modules/generated/sklearn.neural_network.ANNRegressor.html

## References

Anzellotti, S., Caramazza, A., Saxe, R., 2017a. Multivariate pattern dependence. PLoS Comput. Biol. 13, e1005799.

Anzellotti, S., Coutanche, M.N., 2018. Beyond functional connectivity: investigating networks of multivariate representations. Trends Cogn. Sci. 22, 258–269.

Anzellotti, S., Fedorenko, E., Kell, A.J.E., Caramazza, A., Saxe, R., 2017b. Measuring and modeling nonlinear interactions between brain regions with fMRI. bioRxiv 74856.

Basti, A., Mur, M., Kriegeskorte, N., Pizzella, V., Marzetti, L., Hauk, O., 2019. Analysing linear multivariate pattern transformations in neuroimaging data. PLoS One 14.

Basti, A., Nili, H., Hauk, O., Marzetti, L., Henson, R.N., 2020. Multi-dimensional connectivity: a conceptual and mathematical review. Neuroimage 117179.

Basti, A., Pizzella, V., Chella, F., Romani, G.L., Nolte, G., Marzetti, L., 2018. Disclosing large-scale directed functional connections in MEG with the multivariate phase slope index. Neuroimage 175, 161–175.

Bastos, A.M., Schoffelen, J.-M., 2016. A tutorial review of functional connectivity analysis methods and their interpretational pitfalls. Front. Syst. Neurosci. 9, 175.

Bullmore, E., Sporns, O., 2009. Complex brain networks: graph theoretical analysis of structural and functional systems. Nat. Rev. Neurosci. 10, 186–198.

Cabeza, R., 2008. Role of parietal regions in episodic memory retrieval: the dual attentional processes hypothesis. Neuropsychologia 46, 1813–1827.

Cabeza, R., Ciaramelli, E., Moscovitch, M., 2012. Cognitive contributions of the ventral parietal cortex: an integrative theoretical account. Trends Cogn. Sci. 16, 338–352.

Chambers, C.D., Payne, J.M., Stokes, M.G., Mattingley, J.B., 2004. Fast and slow parietal pathways mediate spatial attention. Nat. Neurosci. 7, 217–218.

Chiou, R., Humphreys, G.F., Jung, J., Lambon Ralph, M.A., 2018. Controlled semantic cognition relies upon dynamic and flexible interactions between the executive ‘semantic control’and hub-and-spoke ‘semantic representation’systems. cortex 103, 100–116.

Clarke, A., Devereux, B.J., Randall, B., Tyler, L.K., 2015. Predicting the time course of individual objects with MEG. Cereb. Cortex 25, 3602–3612.

Clarke, A., Taylor, K.I., Tyler, L.K., 2011. The evolution of meaning: spatio-temporal dynamics of visual object recognition. J. Cogn. Neurosci. 23, 1887–1899.

Colclough, G.L., Brookes, M.J., Smith, S.M., Woolrich, M.W., 2015. A symmetric multivariate leakage correction for MEG connectomes. Neuroimage 117, 439–448.

Dhond, R.P., Witzel, T., Dale, A.M., Halgren, E., 2007. Spatiotemporal cortical dynamics underlying abstract and concrete word reading. Hum. Brain Mapp. 28, 355–362.

DiCarlo, J.J., Zoccolan, D., Rust, N.C., 2012. How does the brain solve visual object recognition? Neuron 73, 415–434.

Engemann, D.A., Gramfort, A., 2015. Automated model selection in covariance estimation and spatial whitening of MEG and EEG signals. Neuroimage 108, 328–342.

Ewald, A., Marzetti, L., Zappasodi, F., Meinecke, F.C., Nolte, G., 2012. Estimating true brain connectivity from EEG/MEG data invariant to linear and static transformations in sensor space. Neuroimage 60, 476–488.

Farahibozorg, R., 2018. Uncovering Dynamic Semantic Networks in the Brain Using Novel Approaches for EEG/MEG Connectome Reconstruction. Cambridge.

Farahibozorg, S.-R., 2018. Uncovering Dynamic Semantic Networks in the Brain Using Novel Approaches for EEG/MEG Connectome Reconstruction. University of Cambridge.

Farahibozorg, S.-R., Henson, R.N., Hauk, O., 2018. Adaptive cortical parcellations for source reconstructed EEG/MEG connectomes. Neuroimage 169, 23–45.

Farahibozorg, S.-R., Henson, R.N., Woollams, A.M., Hauk, O., 2022. Distinct Roles for the Anterior Temporal Lobe and Angular Gyrus in the Spatiotemporal Cortical Semantic Network. Cereb. Cortex bhab501. https://doi.org/10.1093/cercor/bhab501

Fries, P., 2015. Rhythms for cognition: communication through coherence. Neuron 88, 220–235.

Glasser, M.F., Coalson, T.S., Robinson, E.C., Hacker, C.D., Harwell, J., Yacoub, E., Ugurbil, K., Andersson, J., Beckmann, C.F., Jenkinson, M., 2016. A multi-modal parcellation of human cerebral cortex. Nature 536, 171–178.

Gramfort, A., Luessi, M., Larson, E., Engemann, D.A., Strohmeier, D., Brodbeck, C., Goj, R., Jas, M., Brooks, T., Parkkonen, L., 2013. MEG and EEG data analysis with MNE-Python. Front. Neurosci. 7, 267.

Gramfort, A., Luessi, M., Larson, E., Engemann, D.A., Strohmeier, D., Brodbeck, C., Parkkonen, L., Hämäläinen, M.S., 2014. MNE software for processing MEG and EEG data. Neuroimage 86, 446–460.

Hagan, M.T., Demuth, H.B., 1999. Neural networks for control, in: Proceedings of the 1999 American Control Conference (Cat. No. 99CH36251). IEEE, pp. 1642–1656.

Hämäläinen, M.S., Ilmoniemi, R.J., 1994. Interpreting magnetic fields of the brain: minimum norm estimates. Med. Biol. Eng. Comput. 32, 35–42.

Hauk, O., 2004. Keep it simple: a case for using classical minimum norm estimation in the analysis of EEG and MEG data. Neuroimage 21, 1612–1621.

Hauk, O., Stenroos, M., Treder, M., 2019. Towards an objective evaluation of EEG/MEG source estimation methods: The Linear Tool Kit. BioRxiv 672956.

Hauk, O., Stenroos, M., Treder, M.S., 2022. Towards an objective evaluation of EEG/MEG source estimation methods–The linear approach. Neuroimage 255, 119177.

Hauk, O., Wakeman, D.G., Henson, R., 2011. Comparison of noise-normalized minimum norm estimates for MEG analysis using multiple resolution metrics. Neuroimage 54, 1966–1974.

Hebb, D.O., 2005. The organization of behavior: A neuropsychological theory. Psychology Press.

Henson, R.N., Mouchlianitis, E., Friston, K.J., 2009. MEG and EEG data fusion: simultaneous localisation of face-evoked responses. Neuroimage 47, 581–589.

Hoerl, A.E., Kennard, R.W., 1970. Ridge regression: Biased estimation for nonorthogonal problems. Technometrics 12, 55–67.

Humphreys, G.F., Lambon Ralph, M.A., 2015. Fusion and fission of cognitive functions in the human parietal cortex. Cereb. Cortex 25, 3547–3560.

Humphreys, G.F., Lambon Ralph, M.A., Simons, J.S., 2021. A unifying account of angular gyrus contributions to episodic and semantic cognition. Trends Neurosci.

Hyvarinen, A., 1999. Fast and robust fixed-point algorithms for independent component analysis. IEEE Trans. Neural Networks 10, 626–634.

Hyvärinen, A., Oja, E., 2000. Independent component analysis: algorithms and applications. Neural networks 13, 411–430.

Jackson, R.L., 2021. The neural correlates of semantic control revisited. Neuroimage 224, 117444.

Jackson, R.L., Hoffman, P., Pobric, G., Lambon Ralph, M.A., 2016. The semantic network at work and rest: differential connectivity of anterior temporal lobe subregions. J. Neurosci. 36, 1490–1501.

Jung, J., Lambon Ralph, M.A., 2016. Mapping the dynamic network interactions underpinning cognition: a cTBS-fMRI study of the flexible adaptive neural system for semantics. Cereb. Cortex 26, 3580–3590.

Jung, J., Rice, G., Lambon Ralph, M.A., 2019. The neural bases of resilient cognitive systems: Evidence of variable neuro-displacement in the semantic system. bioRxiv 716266.

Karimi-Rouzbahani, H., Woolgar, A., Henson, R., Nili, H., 2022. Caveats and nuances of model-based and model-free representational connectivity analysis. Front. Neurosci. 16.

Khalid, S., Khalil, T., Nasreen, S., 2014. A survey of feature selection and feature extraction techniques in machine learning, in: 2014 Science and Information Conference. IEEE, pp. 372–378.

Kietzmann, T.C., Spoerer, C.J., Sörensen, L.K.A., Cichy, R.M., Hauk, O., Kriegeskorte, N., 2019. Recurrence is required to capture the representational dynamics of the human visual system. Proc. Natl. Acad. Sci. 116, 21854–21863.

Kriegeskorte, N., Mur, M., Bandettini, P.A., 2008. Representational similarity analysis-connecting the branches of systems neuroscience. Front. Syst. Neurosci. 2, 4.

Laakso, A., Cottrell, G., 2000. Content and cluster analysis: assessing representational similarity in neural systems. Philos. Psychol. 13, 47–76.

Lambon Ralph, M.A., Jefferies, E., Patterson, K., Rogers, T.T., 2016. The neural and computational bases of semantic cognition. Nat. Rev. Neurosci. 18, 42–55. https://doi.org/10.1038/nrn.2016.150

Liu, A.K., Dale, A.M., Belliveau, J.W., 2002. Monte Carlo simulation studies of EEG and MEG localization accuracy. Hum. Brain Mapp. 16, 47–62.

Maris, E., Oostenveld, R., 2007. Nonparametric statistical testing of EEG-and MEG-data. J. Neurosci. Methods 164, 177–190.

McClelland, J.L., Rumelhart, D.E., 1989. Explorations in parallel distributed processing: A handbook of models, programs, and exercises. MIT press.

McCulloch, W.S., Pitts, W., 1943. A logical calculus of the ideas immanent in nervous activity. Bull. Math. Biophys. 5, 115–133.

Molins, A., Stufflebeam, S.M., Brown, E.N., Hämäläinen, M.S., 2008. Quantification of the benefit from integrating MEG and EEG data in minimum ℓ2-norm estimation. Neuroimage 42, 1069–1077.

Ng, A., 2012. Clustering with the k-means algorithm. Mach. Learn.

Nolte, G., Bai, O., Wheaton, L., Mari, Z., Vorbach, S., Hallett, M., 2004. Identifying true brain interaction from EEG data using the imaginary part of coherency. Clin. Neurophysiol. 115, 2292–2307.

Noonan, K.A., Jefferies, E., Visser, M., Lambon Ralph, M.A., 2012. Aligning evidence from functional neuroimaging and neuropsychology for the neural network underpinning semantic control: A metaanalytic investigation. Manuscr. Submitt. Publ.

Palva, J.M., Wang, S.H., Palva, S., Zhigalov, A., Monto, S., Brookes, M.J., Schoffelen, J.-M., Jerbi, K., 2018. Ghost interactions in MEG/EEG source space: A note of caution on inter-areal coupling measures. Neuroimage 173, 632–643.

Pascual-Marqui, R.D., 2007. Instantaneous and lagged measurements of linear and nonlinear dependence between groups of multivariate time series: frequency decomposition. arXiv Prepr. arXiv0711.1455.

Passingham, R.E., Stephan, K.E., Kötter, R., 2002. The anatomical basis of functional localization in the cortex. Nat. Rev. Neurosci. 3, 606–616.

Rahimi, S., Farahibozorg, S.-R., Jackson, R., Hauk, O., 2022a. Task modulation of spatiotemporal dynamics in semantic brain networks: an EEG/MEG study. Neuroimage 246, 118768.

Rahimi, S., Jackson, R.L., Farahibozorg, S.-R., Hauk, O., 2022b. Time Lagged Multidimensional Pattern Connectivity (TL MDPC): An EEG/MEG Pattern Transformation Based Functional Connectivity Metric. bioRxiv 2022.05.21.492913. https://doi.org/10.1101/2022.05.21.492913

Rice, G.E., Lambon Ralph, M.A., Hoffman, P., 2015. The roles of left versus right anterior temporal lobes in conceptual knowledge: an ALE meta-analysis of 97 functional neuroimaging studies. Cereb. Cortex 25, 4374–4391.

Rogers, T.T., Cox, C.R., Lu, Q., Shimotake, A., Kikuchi, T., Kunieda, T., Miyamoto, S., Takahashi, R., Ikeda, A., Matsumoto, R., 2021. Evidence for a deep, distributed and dynamic code for animacy in human ventral anterior temporal cortex. Elife 10.

Rogers, T.T., McClelland, J.L., 2014. Parallel distributed processing at 25: Further explorations in the microstructure of cognition. Cogn. Sci. 38, 1024–1077.

Rojas, R., 1996. The backpropagation algorithm, in: Neural Networks. Springer, pp. 149–182.

Samuelsson, J.G., Peled, N., Mamashli, F., Ahveninen, J., Hämäläinen, M.S., 2021. Spatial fidelity of MEG/EEG source estimates: A general evaluation approach. Neuroimage 224, 117430.

Sassenhagen, J., Draschkow, D., 2019. Cluster-based permutation tests of MEG/EEG data do not establish significance of effect latency or location. Psychophysiology 56, e13335.

Seth, A.K., 2010. A MATLAB toolbox for Granger causal connectivity analysis. J. Neurosci. Methods 186, 262–273.

Sharma, Sagar, Sharma, Simone, Athaiya, A., 2017. Activation functions in neural networks. Towar. data Sci. 6, 310–316.

Shimamura, A.P., 2011. Episodic retrieval and the cortical binding of relational activity. Cogn. Affect. Behav. Neurosci. 11, 277–291.

Siegel, M., Donner, T.H., Engel, A.K., 2012. Spectral fingerprints of large-scale neuronal interactions. Nat. Rev. Neurosci. 13, 121–134.

Singh, T.N., Kanchan, R., Verma, A.K., Singh, S., 2003. An intelligent approach for prediction of triaxial properties using unconfined uniaxial strength. Min Eng J 5, 12–16.

Sormaz, M., Jefferies, E., Bernhardt, B.C., Karapanagiotidis, T., Mollo, G., Bernasconi, N., Bernasconi, A., Hartley, T., Smallwood, J., 2017. Knowing what from where: Hippocampal connectivity with temporoparietal cortex at rest is linked to individual differences in semantic and topographic memory. Neuroimage 152, 400–410.

Stefaniak, J.D., Geranmayeh, F., Lambon Ralph, M.A., 2022. The multidimensional nature of aphasia recovery post-stroke. Brain.

Stefaniak, J.D., Halai, A.D., Lambon Ralph, M.A., 2020. The neural and neurocomputational bases of recovery from post-stroke aphasia. Nat. Rev. Neurol. 16, 43–55.

Talebi, N., Nasrabadi, A.M., Mohammad-Rezazadeh, I., Coben, R., 2019. nCREANN: Nonlinear Causal Relationship Estimation by Artificial Neural Network; Applied for Autism Connectivity Study. IEEE Trans. Med. Imaging 38, 2883–2890.

Taulu, S., Kajola, M., 2005. Presentation of electromagnetic multichannel data: the signal space separation method. J. Appl. Phys. 97, 124905.

Venkatesan, P., Anitha, S., 2006. Application of a radial basis function neural network for diagnosis of diabetes mellitus. Curr. Sci. 91, 1195–1199.

Vilberg, K.L., Rugg, M.D., 2008. Memory retrieval and the parietal cortex: a review of evidence from a dual-process perspective. Neuropsychologia 46, 1787–1799.

Visser, M., Jefferies, E., Lambon Ralph, M.A., 2010. Semantic processing in the anterior temporal lobes: a meta-analysis of the functional neuroimaging literature. J. Cogn. Neurosci. 22, 1083–1094.

Wagner, A.D., Shannon, B.J., Kahn, I., Buckner, R.L., 2005. Parietal lobe contributions to episodic memory retrieval. Trends Cogn. Sci. 9, 445–453.

